# Host-selected mutations converging on a global regulator drive an adaptive leap by bacteria to symbiosis

**DOI:** 10.1101/067025

**Authors:** M. Sabrina Pankey, Randi L. Foxall, Ian M. Ster, Lauren A. Perry, Brian M. Schuster, Rachel A. Donner, Matthew Coyle, Vaughn S. Cooper, Cheryl A. Whistler

**Author notes:** Correspondence to: Cheryl Whistler. These authors contributed equally to this work (M. Sabrina Pankey, Randi L. Foxall).

## Abstract

Even though eukaryote health relies on beneficial symbionts, host defenses targeting pathogens create substantial obstacles for the establishment of these essential partnerships. To reveal mechanisms of symbiotic adaptation, we experimentally evolved ecologically distinct bioluminescent *Vibrio fischeri* through *Euprymna scolopes* squid light organs. Serial passaging of *V. fischeri* populations through squid hosts produced eight distinct mutations in the *binK* sensor kinase gene that conferred an exceptional selective advantage demonstrated through both empirical and theoretical analysis. Squid-adaptive *binK* alleles promoted colonization and immune evasion behavior which was mediated by symbiotic polysaccharide (Syp). *binK* variation also produced metabolic convergence with native symbionts, and altered quorum sensing and luminescence. Preexisting coordination of symbiosis traits facilitated an efficient solution where altered function of a regulator was the key to unlock multiple colonization barriers. These results identify a genetic basis for microbial adaptability and underscore the importance of hosts as selective agents that shape emergent symbiont populations.

**Impact statement:** Squid selection on non-native *Vibrio fischeri* drove rapid adaptation through convergent mutations of large effect, unmasking preexisting coordinate regulation of symbiosis.

Major subject areas, keywords, and research organism(s)

Major subject area: Genomics and evolutionary biology

## Introduction

Identifying traits under selection by hosts is crucial to understanding the processes governing nascent symbiotic interactions between animals and microbes. The remarkable efficiency by which some bacteria evolve to gain access to novel host niches indicates such adaptability may be an attribute of some bacterial genomes. Adaptation to a new niche, such as a novel host, may involve reconciliation of constraints imposed by genomic content, conflicting regulation, and pleiotropy (Bedhomme et al. 2012; Morley et al. 2015). Given this context, global regulators could serve as effective targets of selection driving adaptive leaps by pathogenic or mutualistic microbes, as long as essential metabolic pathways are both sufficiently insulated from detrimental effects of mutation, and available for integration with accessory functions (Wolfe et al. 2004; Davenport et al. 2015; Jansen et al. 2015). Studies using experimental evolution have often revealed that adaptation can initially proceed through regulatory changes, but few have identified the underlying mechanisms that promote adaptation or linked these processes to natural symbiotic systems (Kawecki et al. 2012; Bedhomme et al. 2012; Morley et al. 2015).

Members of genus *Vibrio*, halophilic bacteria with a broad distribution in marine and brackish environments, have demonstrated a remarkable adaptive capacity to colonize host niches (Nishiguchi 2002; Guerrero-Ferreira and Nishiguchi 2007; Takemura et al. 2014) and as such are important models for understanding the evolution of host association. Bioluminescent *Vibrio fischeri* are common among marine plankton but the species is best known for its mutualistic light organ symbiosis with squids and fish. *V. fischeri* is a well-known model for the study of social quorum-sensing behavior, where communities of bacteria use diffusible pheromone signal molecules to synchronize gene expression in response to cell density (Waters and Bassler 2005; Schuster et al. 2013; Verma and Miyashiro 2013). In *V. fischeri*, quorum sensing occurs through sequential activation by two different pheromone signals: the first signal ‘primes’ sensitive perception of the second signal by reducing the quorum attenuator small RNA Qrr1 and increasing the levels of the LuxR pheromone sensor (Lupp and Ruby 2004; Miyashiro et al. 2010). In turn, when LuxR binds to the pheromone signals it directly activates the expression of the *lux* bioluminescence operon to produce light, which squid use for counter-illumination camouflage during their nocturnal foraging behavior (Lupp et al. 2003; Jones and Nishiguchi 2004). Additionally, the symbiotic association between *V. fischeri* and the squid *Euprymna scolopes* has become a powerful model for interrogating mechanisms underlying bacterial colonization of metazoan host mucosal surfaces where colonists must overcome host defenses that limit infection by non-symbiotic bacteria, including pathogens (Figure 1A). Once newly hatched squid entrap bacteria in mucus near the light organ, symbionts aggregate and then chemotax to pores at the entrance of the nascent light organs (Nyholm et al. 2000). As bacteria swim down the ducts and into the crypts, they face a ‘gauntlet’ of defenses that includes host derived oxidative species (Weis et al. 1996; Small and McFall-Ngai 1999; Davidson et al. 2004) and patrolling macrophage-like haemocytes that exhibit higher affinity in attachment to non-symbionts (Nyholm and McFall-Ngai 1998; Koropatnick and Kimbell 2007; Nyholm et al. 2009). These sanctions ensure that only the correct symbiotic partner gains access to the crypts where host-provided nutrients support bacterial growth (Graf and Ruby 1998; Heath-Heckman and McFall-Ngai 2011). Striking parallels between beneficial *V. fischeri* colonization and pathogenic infection models suggest that the selective pressures exerted by animal hosts may act on a common repertoire of bacterial traits used to circumvent host defensive obstacles (Nyholm and McFall-Ngai 2004).

**Figure 1:**
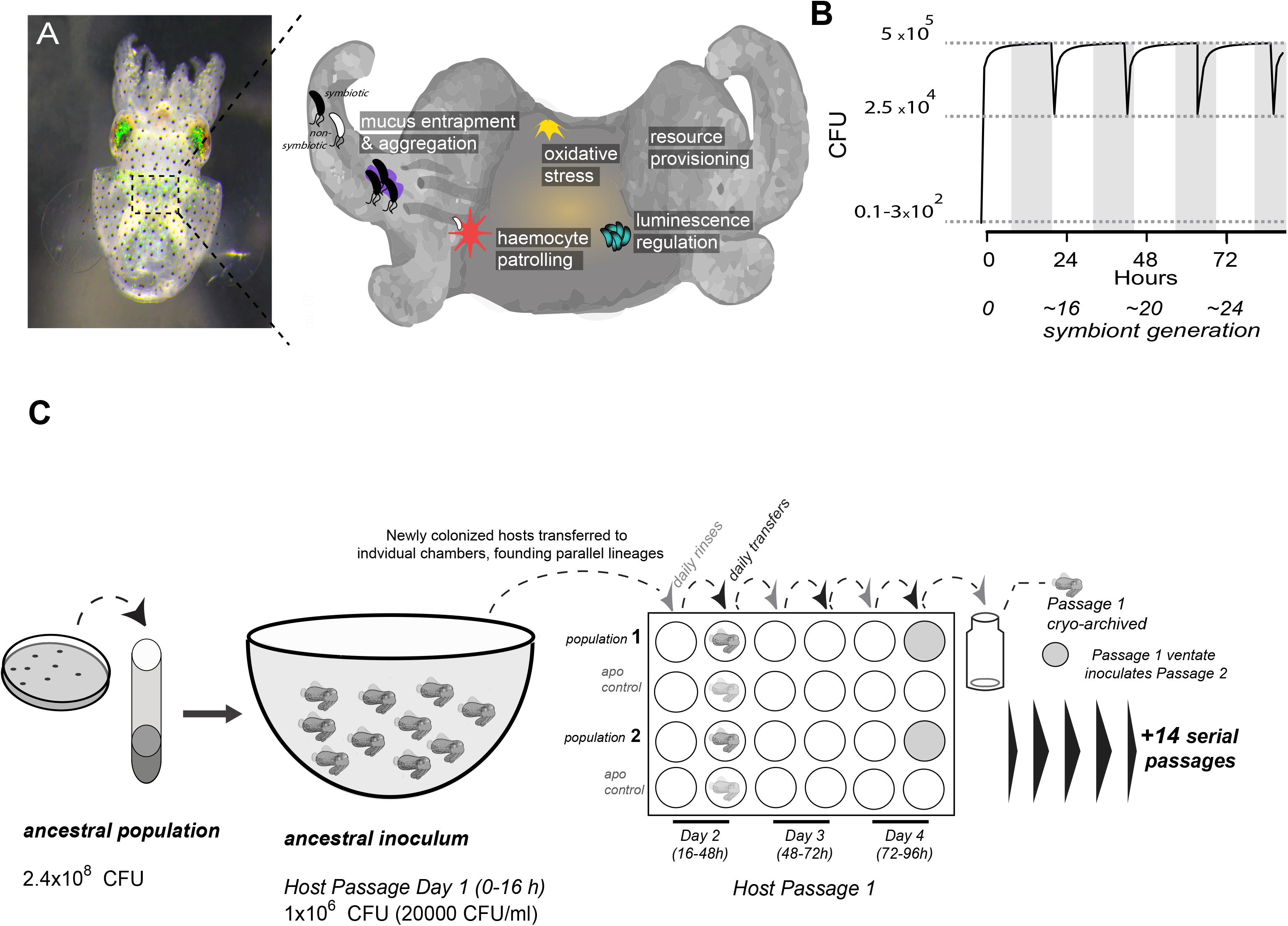
Host selection mechanisms shape symbiotic adaptation in *V. fischeri*. **A)** Host-imposed selection during the temporal process of squid-*V. fischeri* symbiosis from host recruitment (mucus entrapment, aggregation at light organ pores), initiation (host defenses including haemocyte engulfment and oxidative stress), through colonization and maintenance (nutrient provisioning, sanctioning of non-luminous cheaters, and daily purging). **B)** Symbiont population growth modeled for a single passage based on growth dynamics of *V. fischeri* ES114 of initiation population of as few as ~10 (Wollenberg and Ruby 2009; Altura et al. 2013)or as much as 1% inoculum and venting 95% of the population at dawn dawn (every 24 hours) (Boettcher et al. 1996). Shaded areas represent night period whereas light areas represent daylight which induces the venting of 95% of light organ bacteria. **C)** Experimental adaptation *of V. fischeri* symbionts using host selection. Each strain of *V. fischeri* was directly plated from –80°C glycerol stock onto SWT agar plates and grown over-night at 28°C. The bacteria from five colonies were recovered, inoculated into 3 ml SWT broth, and grown until the culture reached mid-log growth (~10^7^ cells per mL), at which time they were diluted into 100 mL seawater at a concentration sufficient to colonize squid based on intrinsic starting capacity (≤20,000 CFU/mL). On day 1, A cohort of ten, 24h-old un-colonized juvenile squid were communally inoculated by overnight incubation, and then moved to individual vials with 4mL FSW where luminescence was monitored. Each squid was then transferred to individual wells of a 24-well polystyrene plate containing filtered sea water with intervening rows of squid from an un-inoculated control cohort (‘apo control’). Note: Only two of the ten passage squid are shown. On days 2, 3, and 4, after venting (which occurred at 36, 60, 84 h post-inoculation), the squid were rinsed by transfer to 2 mL seawater in the adjacent well of the plate, and then transferred once again to the next adjacent column where they were held. Luminescence was measured at various intervals for each squid to monitor colonization and absence of contamination in aposymbiotic control squid. On the fourth day, the squid and half the ventate were frozen at –80°C to preserve bacteria, and the remaining 1 mL ventate was combined with 1 mL of fresh filtered seawater, and used to inoculate a new uncolonized 24-hr old juvenile squid. The process continued for 15 squid only for those lineages where squid were detectably luminous at 48-hours post inoculation.

Not all lineages of *V. fischeri* have attained proficiency in symbiosis, reflective of varied selective regimes that shaped genetic variation and, potentially, adaptability as symbionts (Lee and Ruby 1994a; Nishiguchi et al. 1998). In habitats where they are present, squid hosts control local *V. fischeri* populations by enriching from the planktonic community those strains that are most adept at symbiosis (Lee and Ruby 1994c). Squid recruit small founder populations (~10 bacteria) and subject these to daily cycles of expulsion (‘venting’) and regrowth of 95% of light organ populations to >10^5^ bacteria (Wollenberg and Ruby 2009) (Figure 1A-B) thereby increasing the relative abundance of their light organ inhabitants in the surrounding seawater (Lee and Ruby 1994c). These bottlenecks limit light organ microbial diversity, including variation that impairs symbiosis such as cheater strains that do not contribute to the mutualism (Ruby and McFall-Ngai 1999; Visick and McFall-Ngai 2000; Wollenberg and Ruby 2009). However, host-imposed selection that drives evolution of some lineages towards efficient colonization could hinder future adaptation and entail fitness trade-offs in other environments where symbionts must also persist (Caley and Munday 2003; Soto et al. 2014). By contrast, planktonic *V. fischeri* strains that reside in habitats without hosts, or that are unable to compete for prime host niches, may maintain greater adaptability despite their ineffectiveness as symbionts (Takemura et al. 2014). Deficiency in squid colonization correlates with insufficient or excessive luminescence or inadequate symbiotic polysaccharide (Syp), whose production is controlled by a horizontally-acquired activator (RscS) in squid native strain ES114 (Nishiguchi et al. 1998; Yip et al. 2006; Mandel et al. 2009). However, RscS does not strictly determine squid colonization capacity, indicating other factors contribute to niche breadth. Genomic similarity among closely related yet ecologically diverse strains has obscured relevant functional differences that are sometimes undetectable except in the symbiotic context (Yip et al. 2006; Mandel et al. 2009; Travisano and Shaw 2013).

For this study, we developed a model of experimental evolution in hatchling squid to investigate the genomic underpinnings of partner selection (Schuster et al. 2010). We chose as ancestors of our experimental lineages five *V. fischeri* strains with variable aptitudes for squid symbiosis that were isolated from different niches, including light organs of squid and fish, and planktonic aquatic environments, including one without known hosts (Figure 2A & Table 1). After we experimentally evolved the populations, we evaluated the genetic and phenotypic changes that occurred under host selection to examine how starting fitness and past evolutionary history influence adaptability to squid symbiosis, with the aim of identifying the underlying mechanisms that enable symbiotic adaptation.

**Figure 2:**
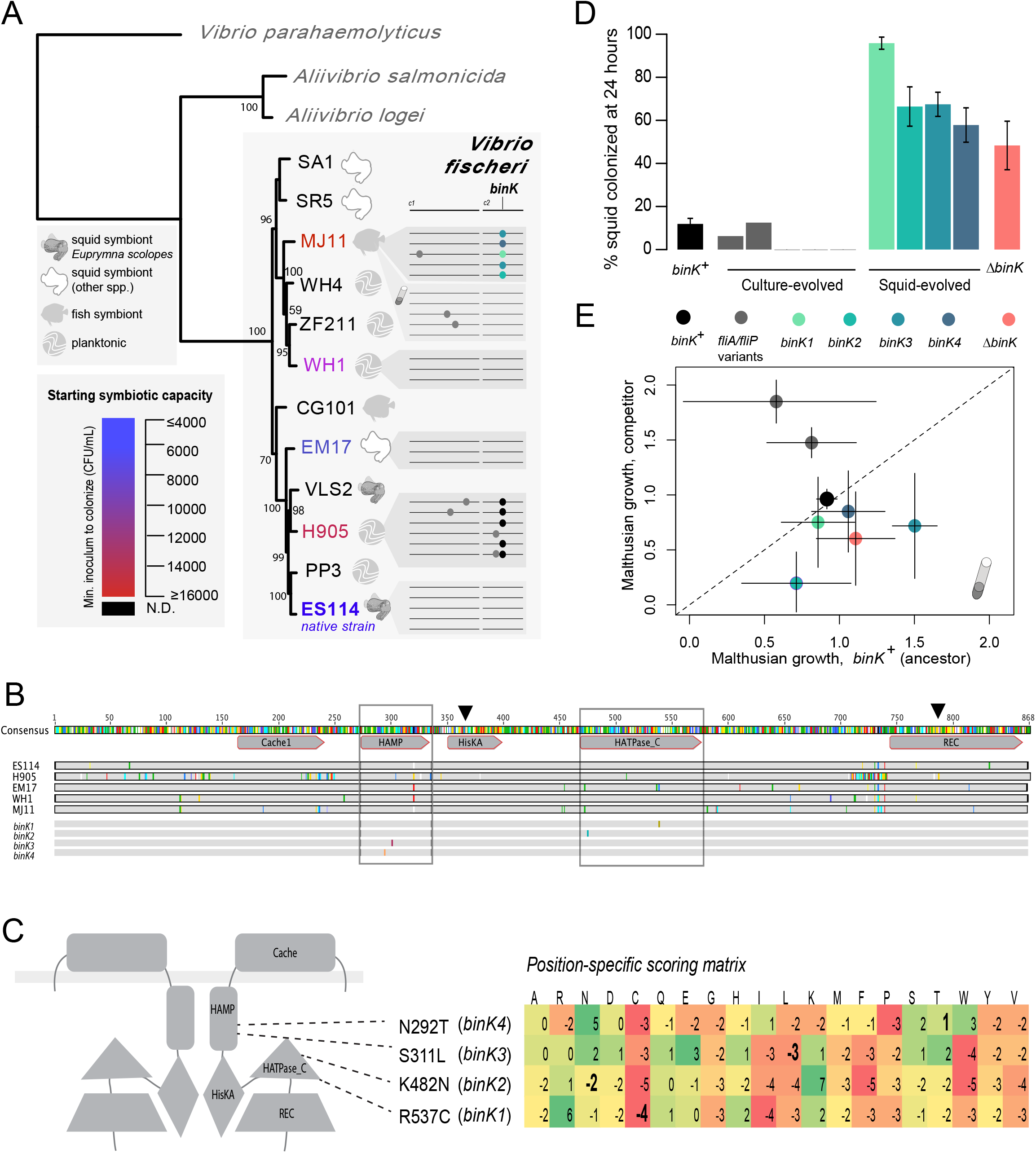
**A)** Phylogenetic relationship, symbiotic capacity, and mutations accrued during squid experimental evolution of ecologically diverse *Vibrio fischeri* strains. Strain relationships were inferred under maximum likelihood using whole genomes with RealPhy (Bertels et al. 2014) and with node supports were calculated from 1000 bootstraps. Graphic symbols for ecological niches represent source of isolation. Intrinsic squid symbiotic capacity for the five experimentally evolved strains as determined by minimum inoculum concentration required for successful initiation of 90% of squid with a 3hr (ES114, EM17, and WH1), or over-night (H905 and MJ11) inoculum represented by color spectrum. At right, consensus genomes for each of the parallel *V. fischeri* populations evolved through *E. scolopes*, with variants indicated by circles. Mutation details available in Tables 1 and 2. Mutations selected in host-passaged populations improved symbiotic capacity not general rigor. **B)** Alignments of the hybrid histidine kinase BinK (locus VFA0360 in strain ES114) and its orthologs in *V. fischeri* strains evolved in this study. Conserved functional domains (Cache1, HAMP, HisKA, HATPaseC and REC) were annotated using *V. fischeri* ES114 RefSeq YP_206318.1. Phosphorylation sites are denoted by arrows. *binK* alleles descended from MJ11 under squid selection are italicized. **C)** BinK mutations arising in squid-evolved populations of MJ11 occurred in the HAMP and HATPase domains. Predicted homodimer structural model based on transmembrane prediction using TMPRed and HMMER and hybrid histidine kinase domain modelling(Anantharaman and Aravind 2000; Stewart and Chen 2010). A position-specific scoring matrix (PSSM) for each of the squid-evolved *binK* sites indicates whether a given amino acid is more (positive) or less (negative) likely to occur than chance. Scores for the substitutions incurred at these sites are shown in bold. **D)** Symbiotic colonization efficiency of MJ11 and derivatives in squid. Percent of squid colonized by culture evolved (c1-c6) and squid evolved (*binK1-remS4*, bolded isolates in Table 2) derivatives of MJ11. Following 3h inoculation of a cohort of 10-20 squid with 3000 CFU/mL of each strain, squid were separated into individual vials, and colonization determined by detectable luminescence at 24 hours. Note: Y-axis is log-scaled. Bars represent standard error. **E)** Growth rates of MJ11 and evolved strains in laboratory culture. Growth of squid native ES114, squid-naïve MJ11, culture-evolved MJ11, 4 lineages of squid-evolved MJ11 and MJ11 *ΔbinK*. Average Malthusian growth rates of *ΔbinK*, squid-evolved *binK* and culture-evolved flagellar mutants (*fliA* and *fliP* variants) following *in vitro* culture competition in minimal media with ancestral *binK^+^* MJ11, estimated using CFU yields of each competitor recovered at regular intervals. Bars: std. err. Diagonal line indicates 1:1 growth.

**Table 1.** Strains and plasmids used in this study.

| **Strain Name** | **Description\*** | **Reference/Source** |
| --- | --- | --- |
| ***Vibrio fischeri* strains<sup>†, ‡</sup>** |  |  |
| ES114 | Isolated from *Euprymna scolopes* | 36 |
| MJ11 | Isolated from *Monocentris japonica* light-organ | 30 |
| EM17 | Isolated from *Euprymna morseii* light-organ | 47 |
| H905 | Isolated from Hawaiian plankton | 38 |
| WH1 | Isolated from Massachusetts plankton | 38 |
| RF1A4 | MJ11 $\Delta binK::ermB$; Em<sup>R</sup> | This study |
| MJ11EP2-3-2 | MJ11 *remS4* | This study |
| MJ11EP2-3-3 | MJ11 *remS4* | This study |
| MJ11EP2-3-4 | MJ11 *remS4* | This study |
| MJ11EP2-3-5 | MJ11 *remS4* | This study |
| MJ11EP2-3-6 | MJ11 *remS4* | This study |
| MJ11EP2-3-7 | MJ11 *remS4* | This study |
| MJ11EP2-3-8 | MJ11 *remS4* | This study |
| MJ11EP15-3-1 | MJ11 *remS4* | This study |
| MJ11EP15-3-3 | MJ11 *remS4* | This study |
| MJ11EP15-3-4 | MJ11 *remS4* | This study |
| MJ11EP15-3-7 | MJ11 *remS4* | This study |
| MJ11EP15-3-8 | MJ11 *remS4* | This study |
| MJ11EP2-4-1 | MJ11 *binK1* | This study |
| MJ11EP2-4-3 | MJ11 *binK1* | This study |
| MJ11EP2-4-4 | MJ11 *binK1* | This study |
| MJ11EP2-4-5 | MJ11 *binK1* | This study |
| MJ11EP2-4-6 | MJ11 *binK1* | This study |
| MJ11EP15-4-1 | MJ11 *binK1 tadC1*<sup>G593T</sup> | 46 |
| MJ11EP15-4-6 | MJ11 *binK1* | This study |
| MJ11EP15-4-7 | MJ11 *binK1* | This study |
| MJ11EP15-4-8 | MJ11 *binK1* | This study |
| MJ11EP2-5-2 | MJ11 *binK3* | This study |
| MJ11EP2-5-3 | MJ11 *binK3* | This study |
| MJ11EP2-5-4 | MJ11 *binK3* | This study |
| MJ11EP2-5-5 | MJ11 *binK3* | This study |
| MJ11EP2-5-6 | MJ11 *binK3* | This study |
| MJ11EP15-5-2 | MJ11 *remS4* | This study |
| MJ11EP15-5-3 | MJ11 *binK3* | This study |
| MJ11EP15-5-4 | MJ11 *binK3* | This study |
| MJ11EP15-5-5 | MJ11 *binK3* | This study |
| MJ11EP2-6-1 | MJ11 *remS2* | This study |
| MJ11EP15-6-1 | MJ11 *remS2* | 46 |
| MJ11EP15-6-2 | MJ11 *remS2* | This study |
| MJ11EP15-6-3 | MJ11 *remS2* | This study |

| Strain Name | Description* | Reference/Source |
| --- | --- | --- |
| MJ11EP15-6-4 | MJ11 *remS2* | This study |
| MJ11EP15-6-5 | MJ11 *remS2* | This study |
| MJ11CE4-1 | MJ11 *fliA*<sup>G80D</sup> | This study |
| MJ11CE5-1 | MJ11 *fliP*<sup>Δ476</sup> | This study |
| ***Escherichia coli* strains** |  |  |
| CC118 *λpir* | Δ(*ara-leu*) *araD* Δ*lac74 galE galK phoA20 thi-1 rpsE rpsB argE recA λpir*; Am<sup>R</sup> | 59 |
| NEB® 10-beta | Δ(*ara-leu*)7697 *araD139 fhuA* Δ*lacX74 galK16 galE15 e14-Φ80dlacZΔM15 recA1 relA1 endA1 nupG rpsL* (Str<sup>R</sup>) *rph spoT1* Δ(*mrr-hsdRMS-mcrBC*) | New England Biolabs, Ipswich, MA |
| TOP10 | F- *mcrA* Δ(*mrr-hsdRMS-mcrBC*) Φ80*lacZΔM15 ΔlacX74 recA1 araD139* Δ(*ara-leu*)7697 *galU galK rpsL* (Str<sup>R</sup>) *endA1 nupG* | Invitrogen, Carlsbad, CA |
| **Plasmids** |  |  |
| pCR™2.1-TOPO® | Commercial cloning vector; Am<sup>R</sup> Kn<sup>R</sup> | Invitrogen, Carlsbad, CA |
| pVSV105 | Mobilizable vector; Ch<sup>R</sup> | 58 |
| pRAD2E1 | pVSV105 carrying wild-type *binK*; Ch<sup>R</sup> | This study |
| pRF2A2 | pVSV105 carrying *binK1*; Ch<sup>R</sup> | This study |
| pCLD48 | pVSV105 carrying ES114 *sypE*; Ch<sup>R</sup> | 60 |
| pTM268 | pVSV105 carrying ES114 *P<sub>qrr1</sub>-gfp* and *P<sub>tetA</sub>-mCherry*; Ch<sup>R</sup> | 23 |
| pVSV104 | Mobilizable vector; Kn<sup>R</sup> | 58 |
| pRF2A1 | pVSV104 carrying *sypE*; Kn<sup>R</sup> | This study |
| pKV111 | Mobilizable vector containing *gfp*; Ch<sup>R</sup> | 36 |
| pVSV103 | Mobilizable vector containing *lacZ*; Kn<sup>R</sup> | 58 |
| pCAW7B1 | pVSV103 containing *lacZΔ147-1080bp*; Kn<sup>R</sup> | This study |

\* Em<sup>R</sup>, erythromycin resistance; Kn<sup>R</sup>, kanamycin resistance; Am<sup>R</sup>, ampicillin resistance; Ch<sup>R</sup>, chloramphenicol resistance.
† Experimentally evolved strains are designated 'MJ11EP#-#-#', where the first and second numbers after the 'P' designates the squid passage and population from which the strain was isolated, and the third number designates isolate number; strains derived from evolution in culture designated by 'MJ11CE'.
‡ The complete genome of *Vibrio fischeri* strains in grey text have not been sequenced.

## Results

### *Squid experimental evolution of ecologically diverse* V. fischeri *repeatedly produced adaptive mutations in a sensor kinase gene*

To study the dynamic process of adaptation during symbiosis, we capitalized upon the squid’s natural recruitment process to found parallel populations of *V. fischeri*, and used the daily squid venting behavior to restrict and re-grow bacterial populations, which were passaged through 15 serial squid, encompassing 60 bottlenecking events and an estimated 290-360 generations (Figure 1C). Multiple populations were derived in parallel from each of five ancestral strains using high density inocula up to 10 times the concentration required for native strain colonization, in order to overcome the colonization deficiencies of squid maladapted strains (Figure 2A and Methods).

Genome sequencing of representative evolved isolates revealed few mutations (Figure 2A, Table 2). Among these were several that converged in a hybrid histidine sensor kinase-encoding gene (locus VF_A0360 in *V. fischeri* ES114)(Figure 2B), which was recently identified as a <u>b</u>iofilm <u>i</u>nhibition <u>k</u>inase (*binK*) (Brooks and Mandel 2016). Nine independent mutations mapping to the *binK* locus, most often without other co-occurring mutations, dominated multiple parallel evolved populations of the two strains initially most impaired at squid symbiosis: MJ11 and H905 (Figure 2A and Table 2). Given that MJ11 is a fish symbiont that lacks *rscS*, and H905 is a planktonic isolate from the squid habitat which is a poor squid colonizer despite harboring *rscS*, starting fitness better predicted the path of evolution than *rscS* content or past evolutionary history as inferred by either lineage or lifestyle (Figure 2A)(Lee and Ruby 1994b; Mandel et al. 2009). In contrast, very few mutations arose in lineages with relatively greater starting fitness, and no mutations arose in the representative isolates from the native squid symbiont ES114 (Figure 2A and Table 2). The repeated sweeps of novel *binK* mutations during squid evolution but not during laboratory culture evolution that mimicked the population dynamics of squid-induced bottlenecks suggested that *binK* variants are squid-adaptive rather than the result of mutational hotspots (Table 4)(Dillon et al. 2015).

**Table 2.**
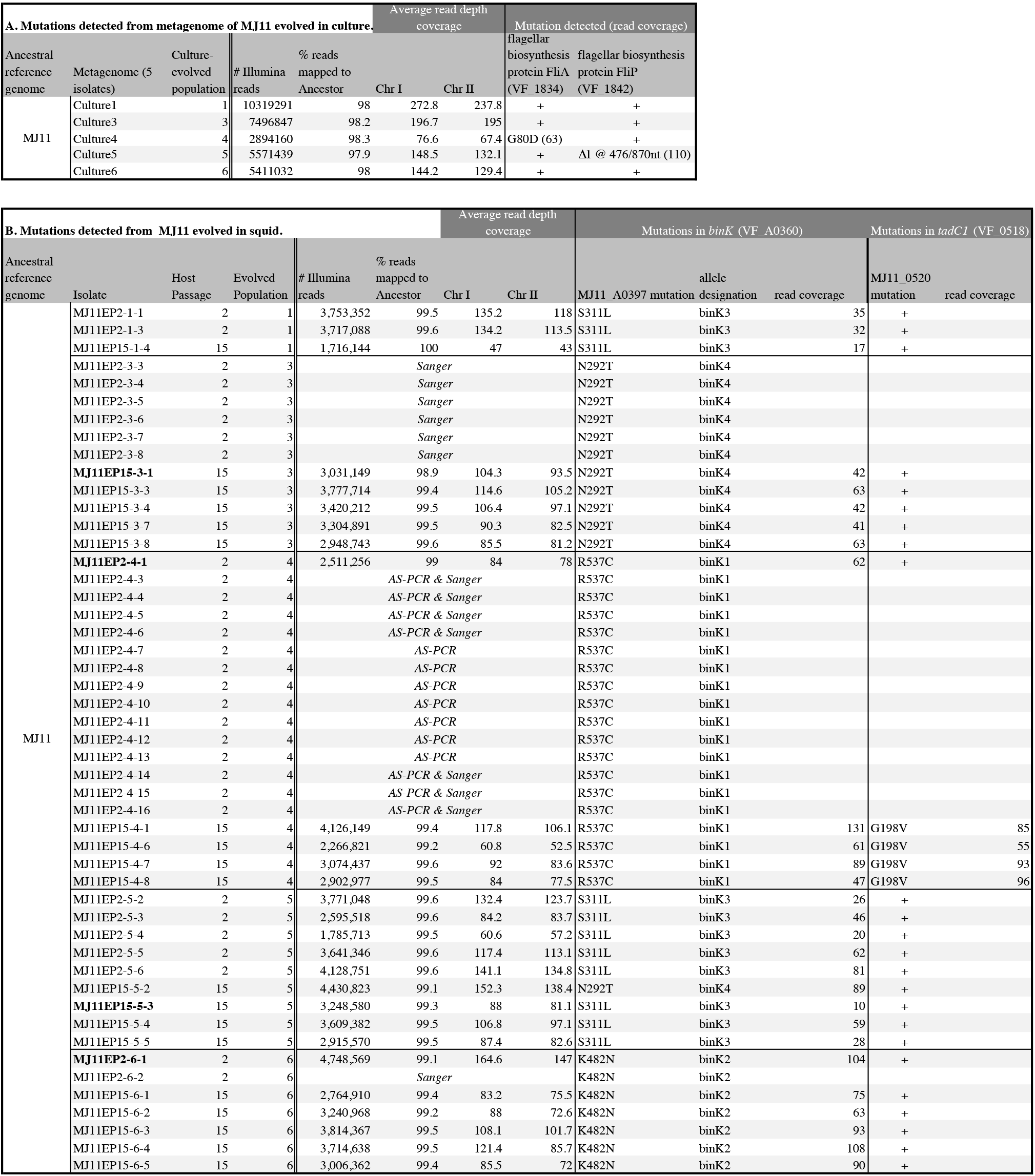
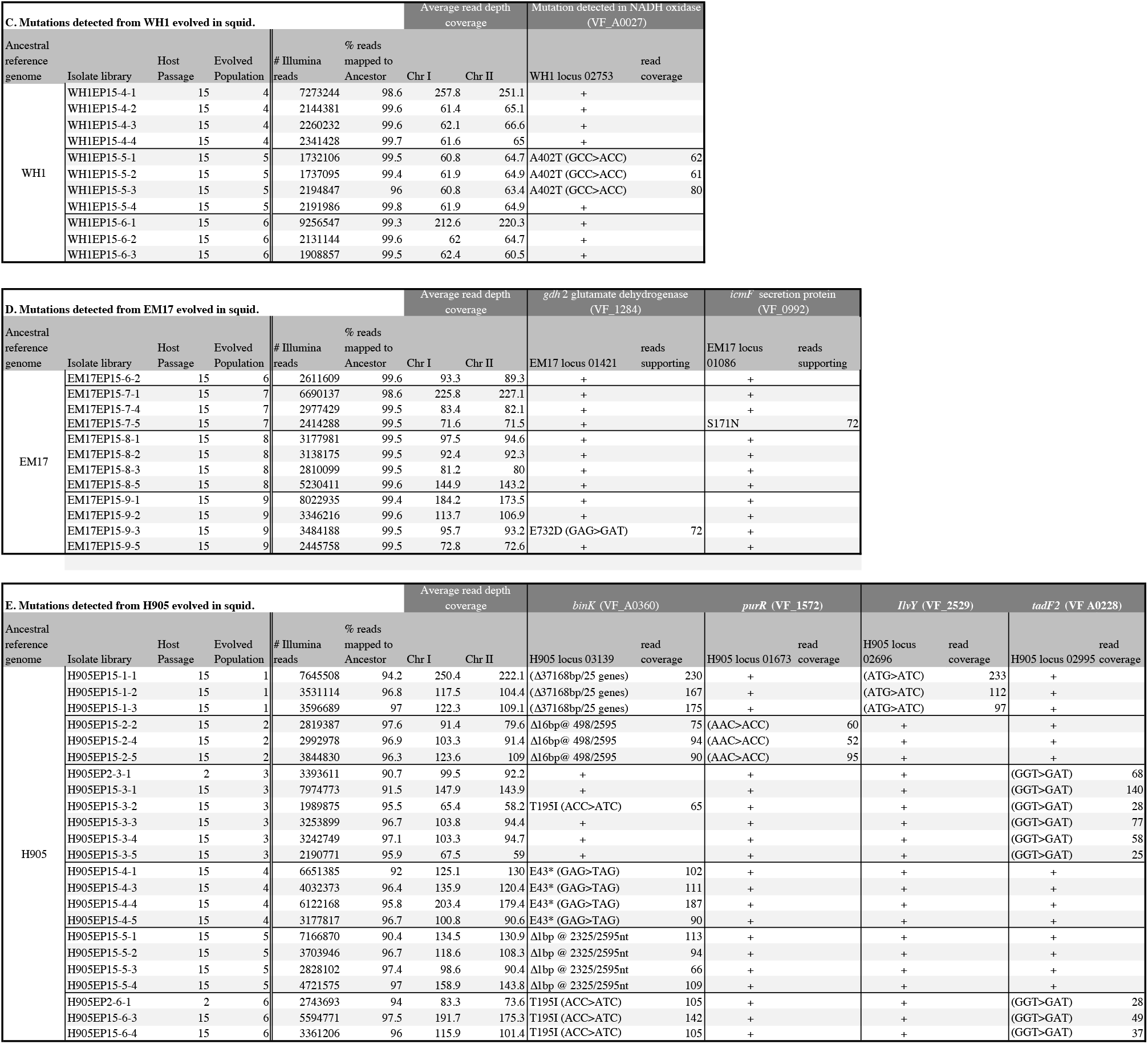
Mutations detected following experimental evolution of *V. fischeri* using Illumina genome resequencing and targeted Sanger sequencing. For culture-evolved populations of *V. fischeri* MJ11, 5 isolates from each evolved population were combined to generate 5 metagenomes. For squid-evolved populations of MJ11, EM17, WH1 and H905, individual isolates were sequenced from lineages that ultimately survived 15 host passages. Mutations were called for sites with minimum read coverage of 20. Isolates saved from early evolutionary time-points (host passage 2) are shown along with isolate genomes from the endpoint (host passage 15). Isolates whose genotypes were confirmed only through allele-specific PCR or Sanger sequencing of the *binK* locus. Coding genes reference *V. fischeri* ES114 locus tags. Isolates in bold served as allelic *binK* representatives for further assays. Mean read depth and coverage of variant sites identified by breseq(Deatherage and Barrick 2014) is also provided.

**Table 3.**
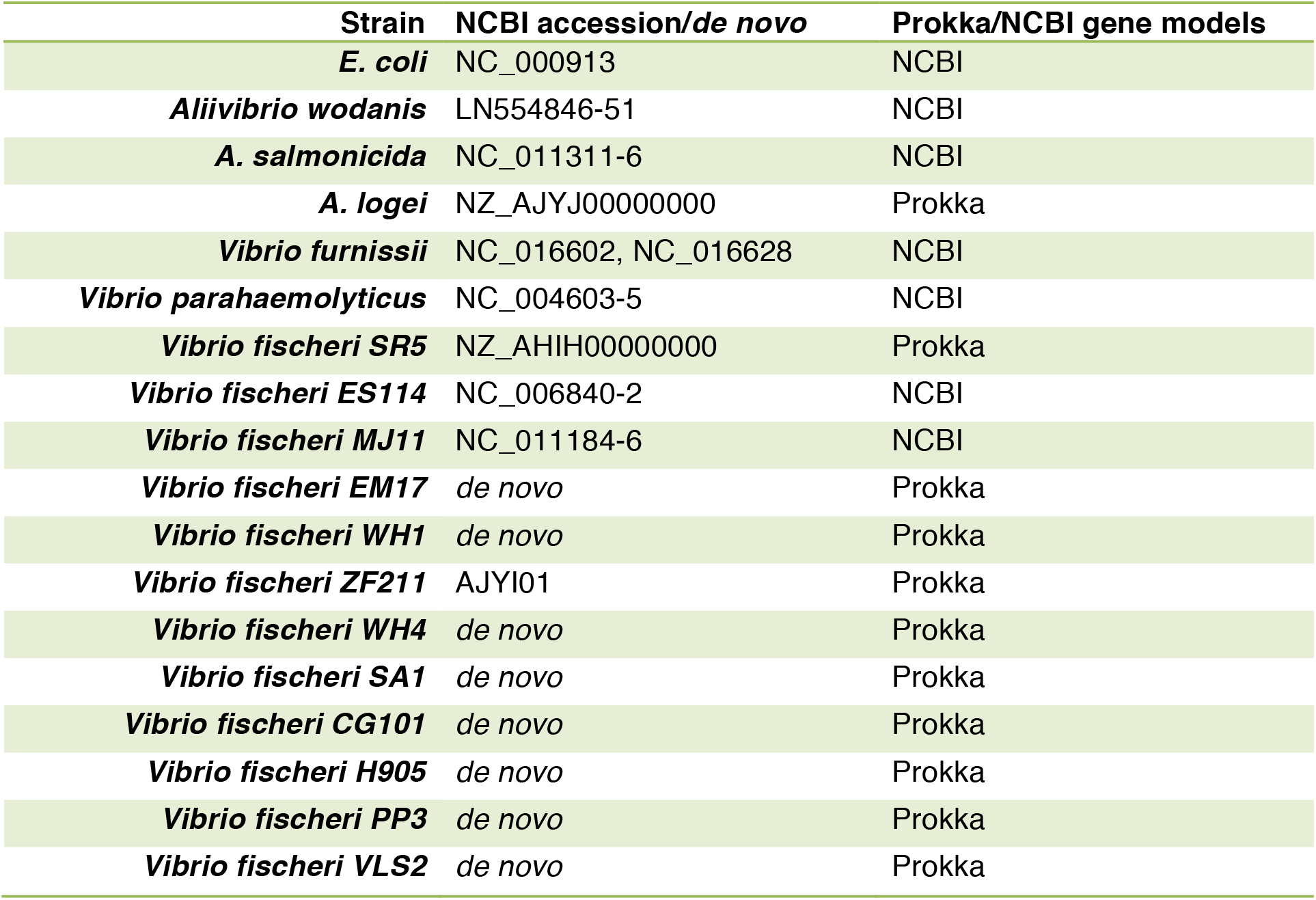
Genomes used in phylogenetic analyses. Provided are GenBank accessions for nucleotide genomes used in strain phylogeny and source for gene models used in hybrid histidine kinase phylogeny.

**Table 4.**
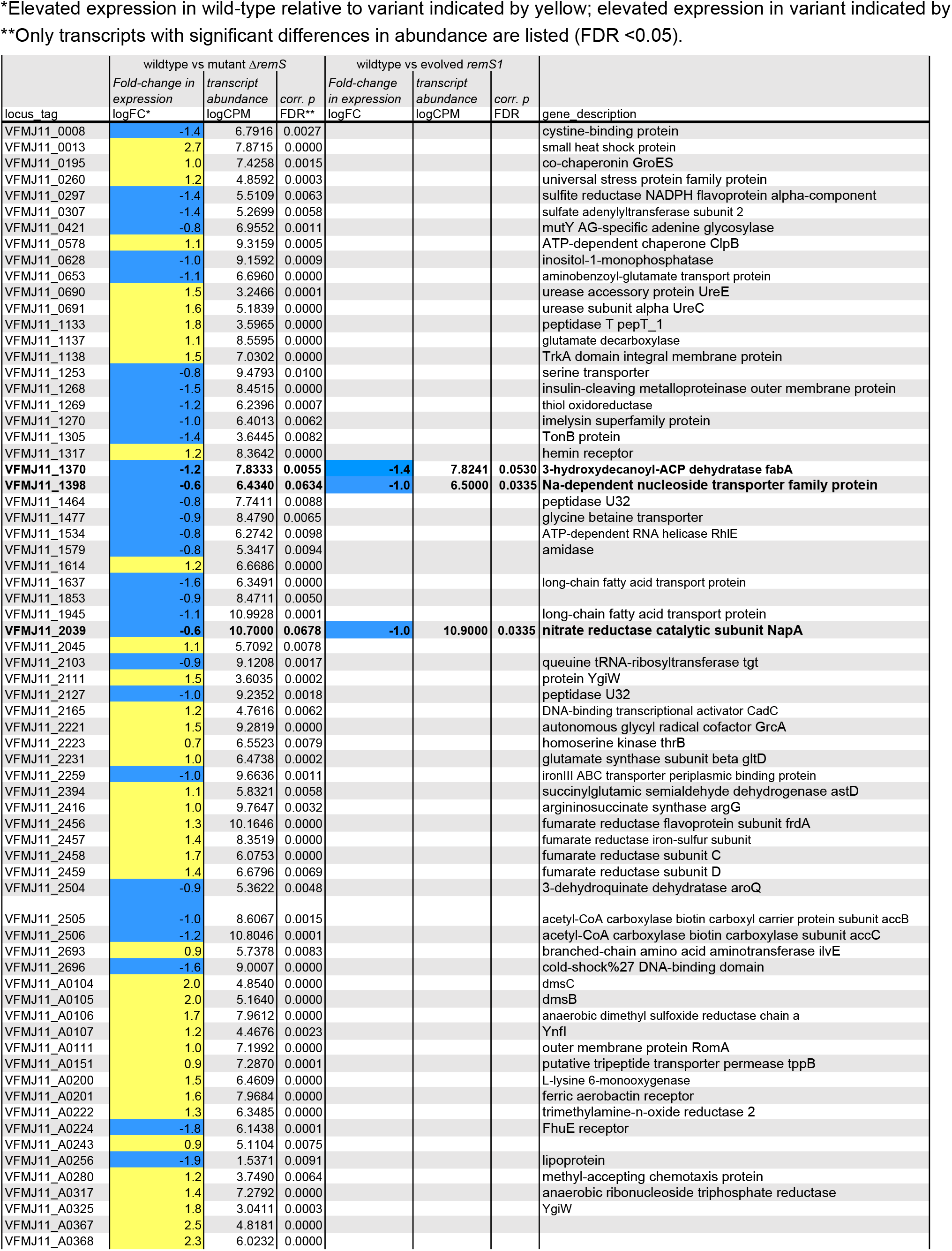
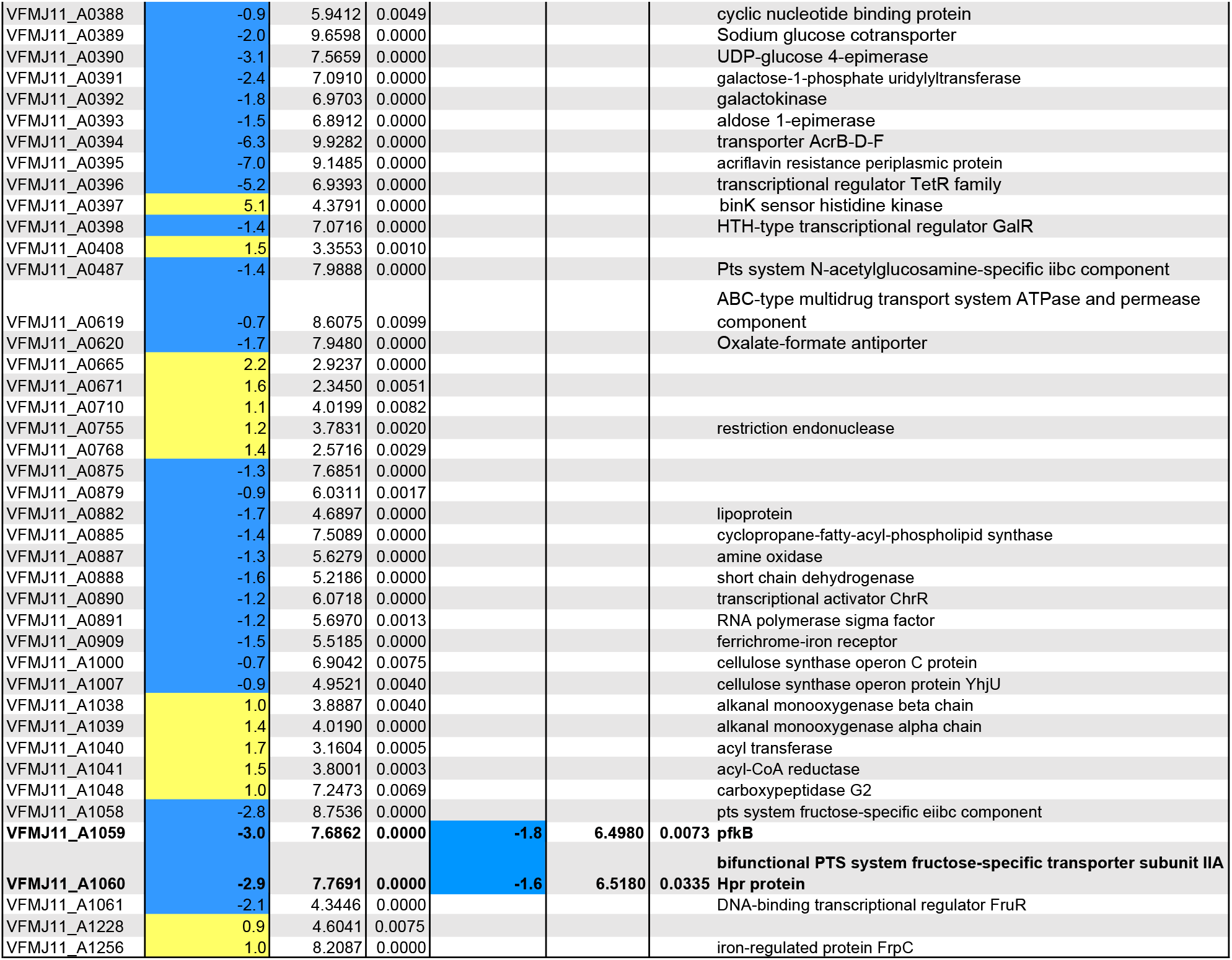
Transcript expression differences between wild-type *binK^+^* (ancestral MJ11) and *binK* mutants (*ΔbinK* and *binK1*) as detected by RNAseq under Fisher’s Exact test in edgeR. ^a^Yellow indicates elevated expression in wild-type relative to mutant indicated; blue indicates reduced expression in wild-type relative to mutants. Loci with similar and significant expression changes in both ΔbinK and *binK1* relative to wild-type listed in bold. ^b^Only loci with significant differences in transcript abundance are listed (FDR <0.05).

Focusing on the evolution of the relatively well-characterized fish symbiont MJ11, only five of ten squid colonized with the same founder population successfully passaged symbionts to the second recipient squid, and each of these populations harbored *binK* mutants (Table 2). Among these were four unique alleles wherein the acquired substitutions mapped to two of the five predicted functional domains of the deduced BinK protein (Figure 2C and Table 2). Analysis of these domains and the acquired substitutions using a position-specific scoring matrix (Figure 2C) indicates each would likely influence function. In each of the five successful squid-evolved lineages of MJ11, *binK* variants dominated the light organ populations by the third experimental squid (Table 4). If beneficial variants in this or any other locus were among the remaining five light organ populations, their failure to colonize the second experimental squid amounted to extinction for these lineages.

### *Squid-adapted* binK *improves fitness during symbiosis initiation and persistent colonization, consistent with theoretical predictions*

Considering convergent evolution is strongly suggestive that the *binK* alleles are adaptive for squid colonization, we directly evaluated the contribution of evolved *binK* alleles to symbiotic colonization. In experiments using inoculum doses typically applied for the native symbiont strain ES114, each squid-evolved *binK* mutant improved colonization efficiency, but were significantly more fit in laboratory culture (indicative of mutants enhancing general vigor) compared to ancestral MJ11 (Figure 2D-E). Moreover, none of five culture-evolved populations of MJ11 accrued *binK* mutations (Table 4) or improved as squid symbionts (Figure 2D). Although mutations mapping to either *binK* domain vastly improved squid colonization relative to the *binK^+^* allele (Figure 2D), variants harboring mutations in of either domain were competitively indistinguishable (t-test, p = 0.34) (Figure 2 supplemental)

To empirically quantify the selective advantage (selective coefficient: *s*) of a *binK* variant (hereafter *binK1* encoding a R537C substitution) harboring no other mutations (Table 2) we coinoculated squid with MJ11 and low densities of *binK1*, simulating the conditions under which these alleles evolved (Figure 3). The estimated selective advantage based on the ratios of the growth rates of each ancestor and evolved variant in co-colonized squid was independent of initial allele frequencies, consistent with a model of hard selection (Figure 3) (Saccheri and Hanski 2006). The estimated selective advantage of the evolved allele nearly doubled between 24 and 48 hours in squid (24 h: 1.1; 48h: 1.8), demonstrating that the competitive advantage extended beyond initiation and initial growth to include the period of re-growth in light organs following expulsion (Figure 3B). In contrast, the squid-evolved *binK* alleles reduced fitness relative to wild-type (*binK^+^*) in laboratory culture (–0.18 > *s* > –1), demonstrating a fitness cost in the absence of hosts (Figure 2E).

**Figure 3.**
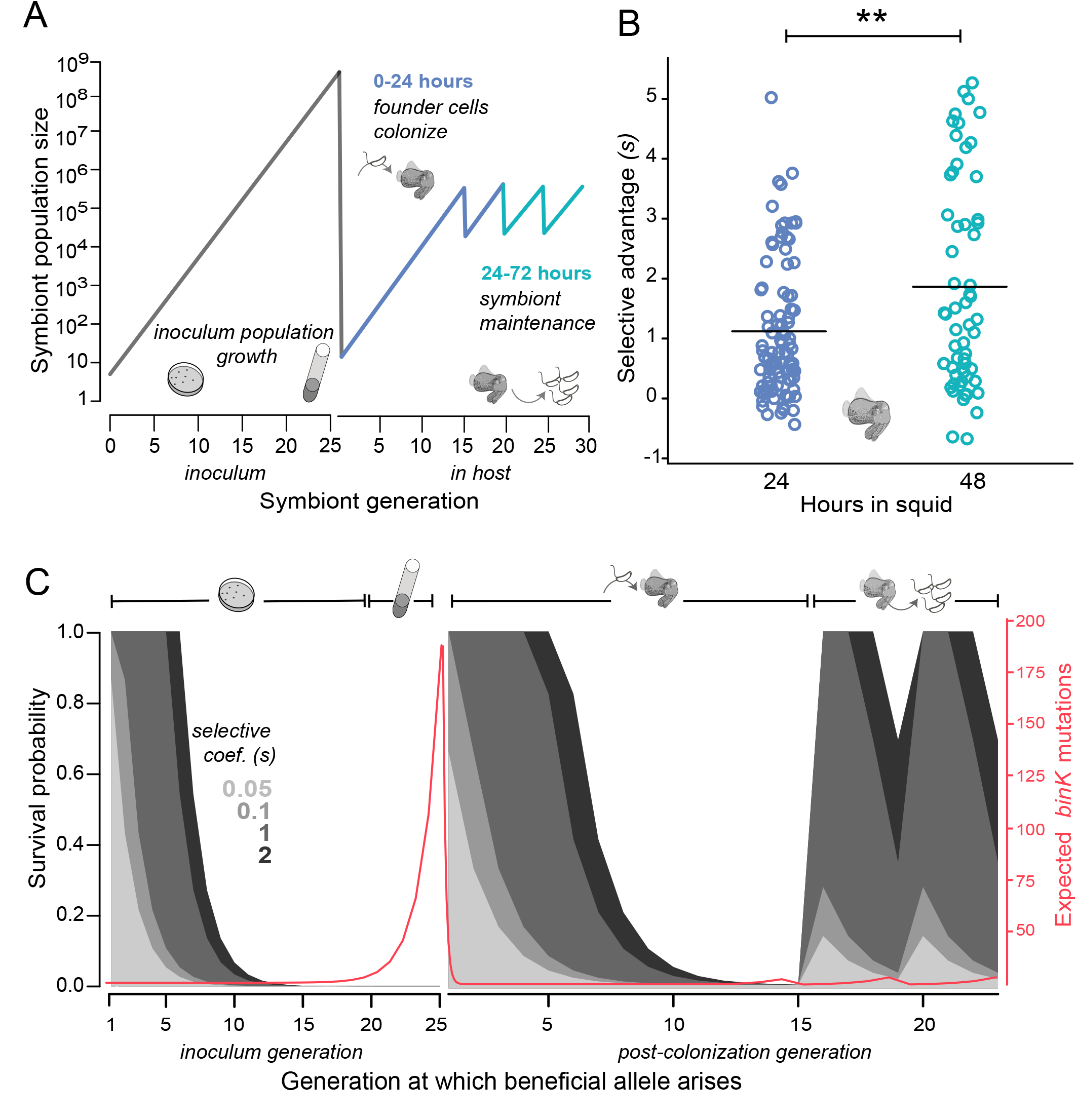
Empirical and modeled estimates of selective advantage in evolving *V. fischeri* symbiont populations. **A)** Symbiont population dynamics during growth in inoculum and following host colonization (black line), including daily host-imposed bottlenecks. **B)** Comparison of selection coefficients conferred by *binK1* in strain MJ11EP2-4-1 (harboring no other mutations) relative to *binK+* from co-inoculated squid light organs after 24 or 48 hours. Selective advantage of the evolved allele increased significantly during this period from 1.1 to 1.8 (Fisher-Pitman Permutation test, **p=0.00108). **C)** Modeled survival probabilities of new beneficial alleles arising in a growing symbiont population facing host-imposed bottlenecks. Red line indicates the number of non-synonymous mutations predicted to accrue within the *binK* locus under neutral evolution during population growth (see Methods). Gray shaded curves model the survival probability of new mutants following the subsequent population bottleneck, which depends on both the generation of growth in inoculum or host in which they arise (x-axis) and the selective advantage s conferred by mutation (gray shading). Notably, beneficial variants arising early in inoculum culture are likely to survive extinction at the subsequent bottleneck, and this probability of survival rapidly decreases even when conferring a large selective coefficient. Based on this model, mutations conferring a large selective advantage (s~1) would have a 50% chance of surviving the subsequent colonization bottleneck if they arose within the first ~6 generations of inoculum growth.

Even with the extreme fitness advantage attained by evolved *binK* variants growing within squid, their repeated recruitment among the few cells that initiated symbiosis is remarkable (Nyholm and McFall-Ngai 2004; Wollenberg and Ruby 2009). Therefore, we modeled the dynamic of evolution over a theoretical range of selection coefficients using population parameters previously documented in the squid-*Vibrio* symbiosis (Wahl and Gerrish 2001; Wollenberg and Ruby 2009; Altura et al. 2013) (see methods) (Figure 3C). The model predicts that in order for beneficial variation to escape extinction during the host-imposed bottleneck during initial colonization, mutants would have to arise early during population expansion and confer s ~ 6. Conversely, any beneficial variants arising in light organs during the maintenance of symbiosis, characterized by daily venting bottlenecks, have increased survival odds even if they confer a lower selective advantage, but the probability of their occurrence is reduced owing to the small effective population size (Methods and Figure 3C). Taking into account the mutation rate determined empirically from mutation accumulation studies with *V. fischeri* ES114 (Sung et al. 2016), even with the relatively few generations that occurred during growth of the inoculum (Figure 3A), we predict as many as 185 *binK* variants could have arisen and been available for host selection under neutral evolution during inoculum growth (Figure 3C). Thus, the empirical estimates of the selective advantage conferred by *binK1* in the symbiotic environment are supported by theoretical estimates from a model of strong selection that promotes beneficial allele survival and enrichment during repeated bottlenecks (Wahl and Gerrish 2001)

### *Host-adapted* binK *improved early colonization behavior and evasion of host immunity through enhanced extracellular matrix and Syp polysaccharide*

We next evaluated which traits conferred the fitness gain by a squid-adaptive *binK* variant during the initiation stage of symbiosis (Figures 1 and 4) (Nyholm and McFall-Ngai 2004). *binK1* improved aggregation at the entrance to the light organs and increased biofilm production compared to MJ11, both of which are traits associated with the Syp symbiotic polysaccharide (Figure 5). Furthermore, transcriptional profiling indicated wild-type *binK^+^* repressed cellulose synthesis, a constituent of biofilm which is co-regulated with Syp (Figures 4 & 5A, Table 4), suggesting the effects of *binK1* on biofilm could be mediated by either or both substrates (Darnell et al. 2008; Visick 2009; Bassis and Visick 2010). Concurrently, the *binK1* variant also resisted binding by host haemocytes to a level comparable to squid-native strain ES114, and survived oxidation better than MJ11 (Figure 4). To evaluate whether Syp polysaccharide mediated these abilities, we over-expressed a repressor of Syp, *sypE* (Morris and Visick 2012) which abolished biofilm formation, haemocyte evasion, and oxidative survival in the *binK1* variant.

**Figure 4.**
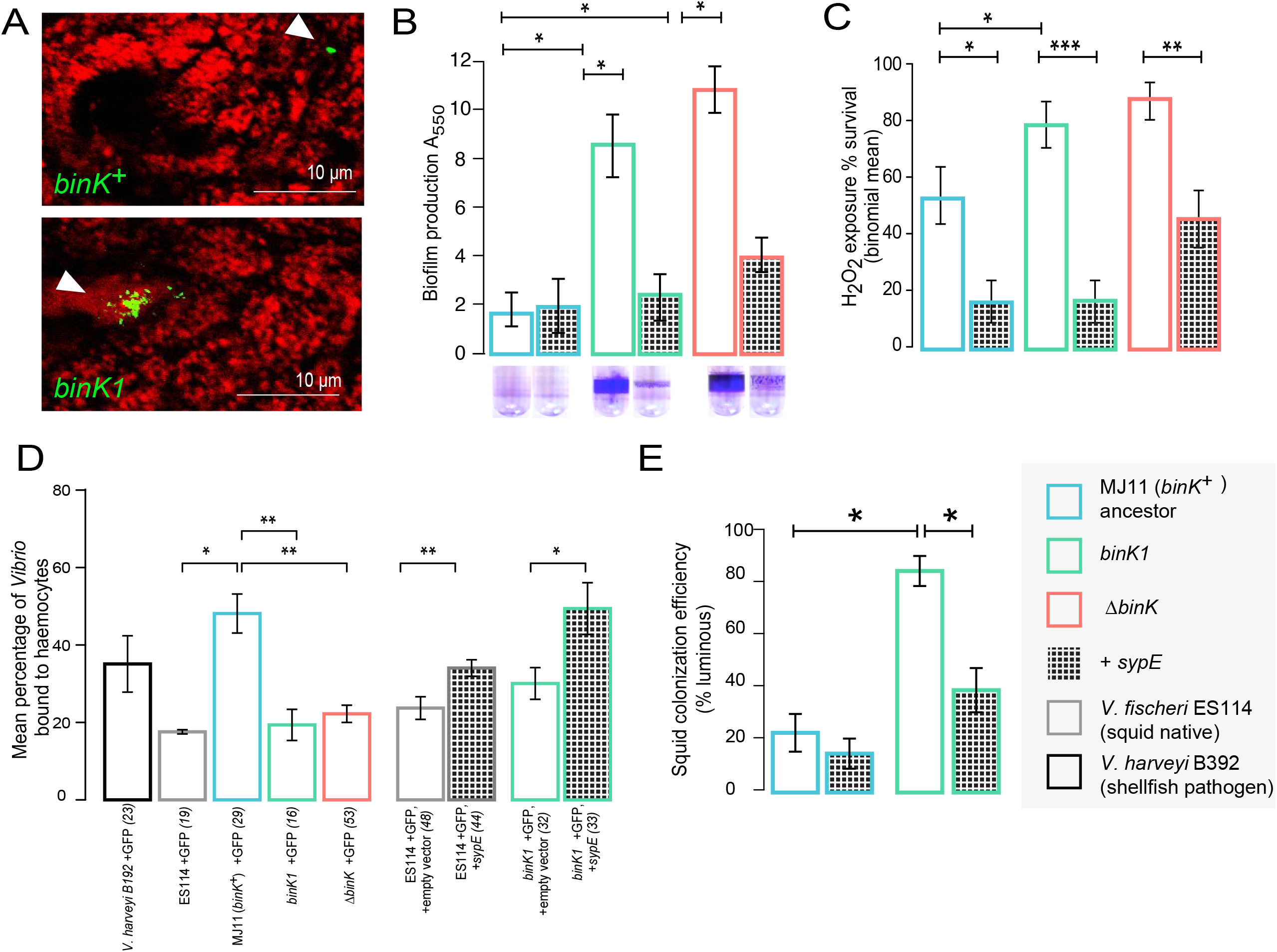
Host-adapted *binK1* improves symbiotic traits through Syp-biofilm. **A)** *V. fischeri* MJ11 aggregate formation near light organ ducts. Host tissue stained with CellTracker Orange. Symbionts carry GFP plasmids (pKV111)(Nyholm et al. 2000). Micrographs show representative *V. fischeri* aggregates following dissection of 30 newly hatched animals incubated with each strain. Aggregates were visualized between 2 and 3 hours of inoculation using a Zeiss LSM 510 Meta laser scanning confocal microscope. **B)** Biofilm production, including in the presence of the Syp-specific repressor *sypE* (pCLD48) or empty vector (pVSV105). **C)** Population survival following exposure to hydrogen peroxide, including in the presence of *sypE* (pCLD48) or control vector (pVSV105). **D)** Evasion of host haemocyte binding by GFP-tagged strains *binK^+^* MJ11, *ΔbinK* MJ11 (RF1A4), and strains ES114 and *binK1* MJ11 carrying either *sypE* (pRF2A1) or control vector (pVSV104). **E)** Colonization efficiency (% colonized squid at 24h) by *binK^+^* and *binK1* strains carrying either *sypE* (pCLD48) or empty vector (pVSV105). Error bars: std. err. *P<0.05, **P<0.01, ***P<0.001.

### *Squid-adapted* binK *confers metabolic convergence with native symbionts*

Transcriptomic and metabolic profiles of *binK* variants suggested that *binK1* also mediated response to substrates related to biofilm production and those supplied by the squid host or important to squid association (Tables 4 & 5, Figure 5). Although transcriptional profiling captured relatively few significant changes in gene expression in the *binK1* variant grown in broth culture, data obtained with the *ΔbinK* mutant suggested that in addition to repressing cellulose synthesis, the wild-type gene also represses carbohydrate glycosylation and sugar transport and metabolism. Notably, the *ΔbinK* mutant increased transcription of serine and N-acetyl-glucosamine transporters. Transcriptional differences also indicated a significant effect of *binK* on iron metabolism and fatty acid biosynthesis pathways associated with luminescence regulation, both of which are important during persistent light organ colonization (Graf and Ruby 1998; Visick et al. 2000; Whitehead et al. 2001; Septer et al. 2011; 2013). However, siderophore production remained undetectable in *binK* variants as it is in the MJ11 ancestor (Supplemental Figure 5).

**Figure 5.**
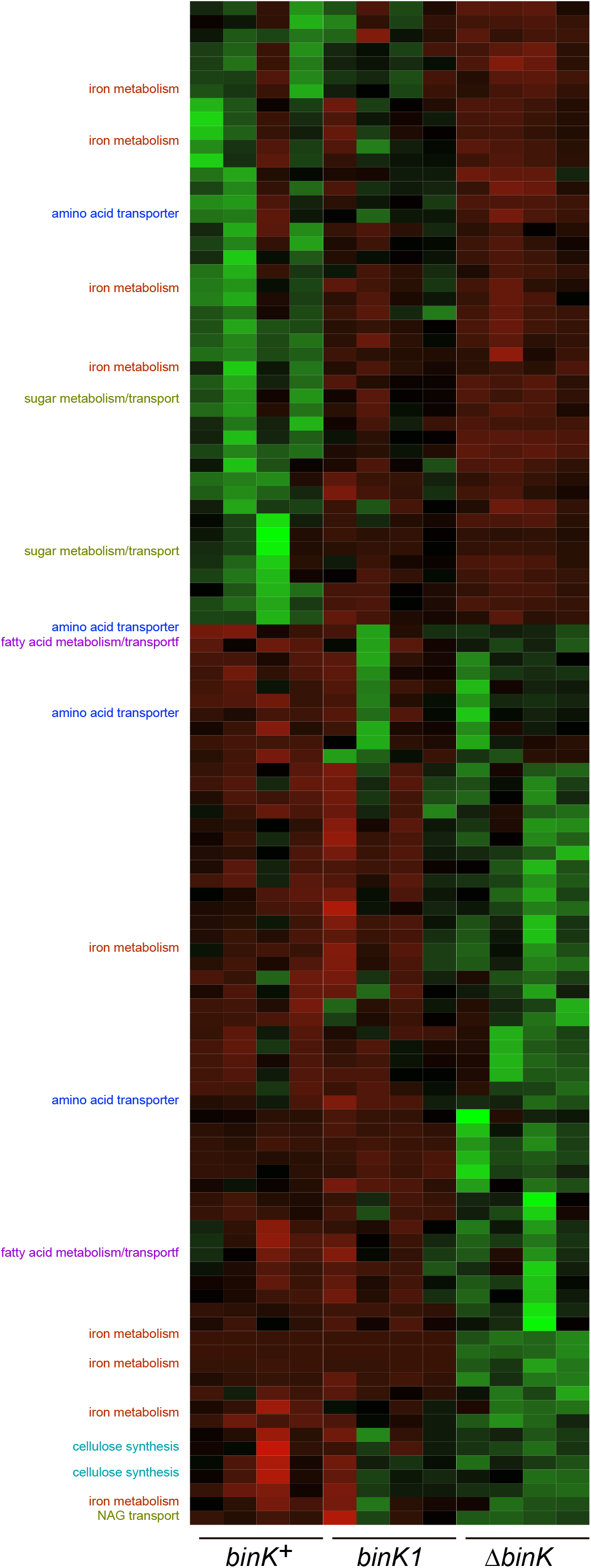
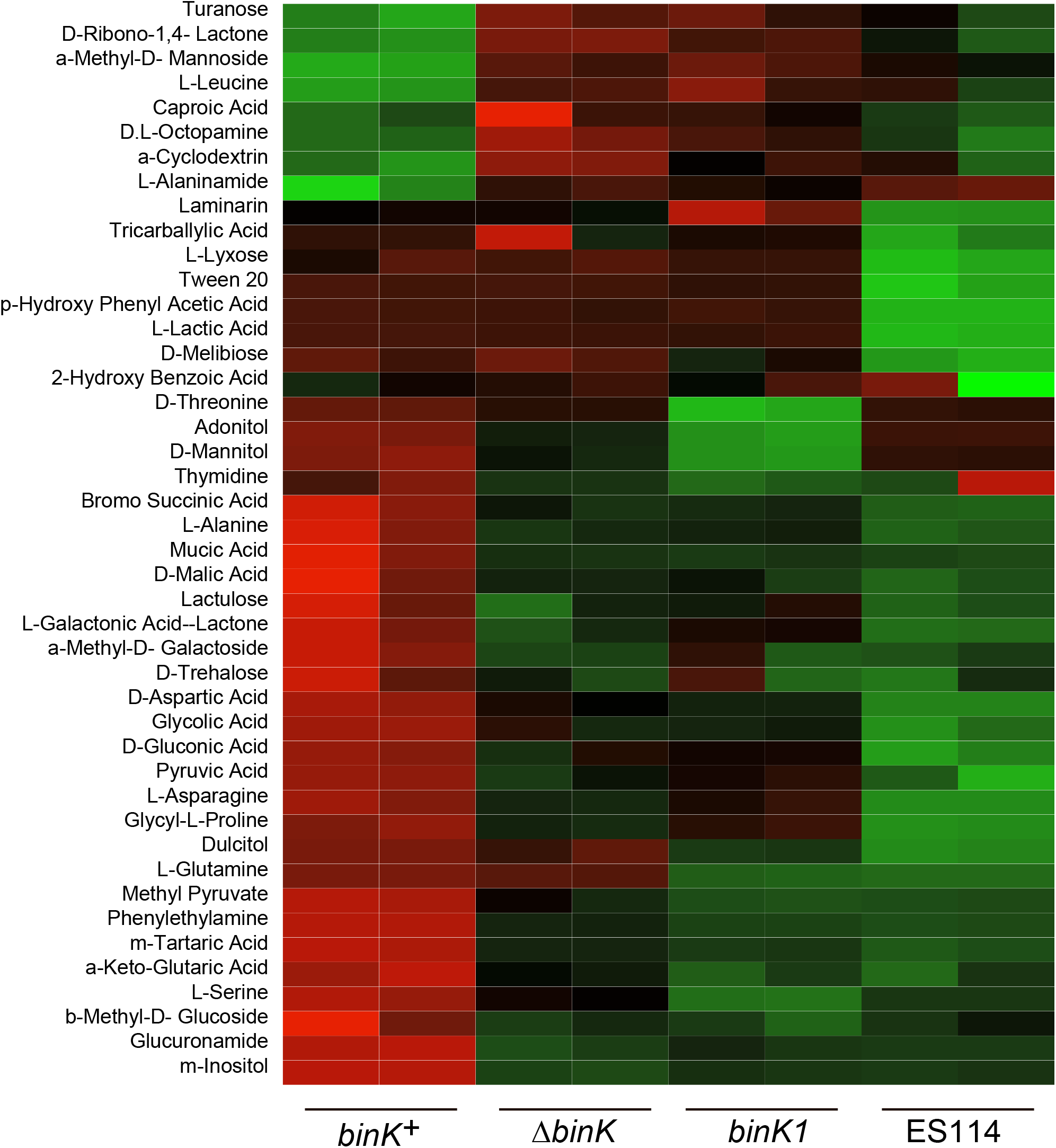
Physiological shifts associated with *binK* variants. **A)** Transcriptomic differences between wild-type MJ11 (*binK^+^*), squid-adapted MJ11 *binK1 and* MJ11 *ΔbinK* for the 3839 coding loci in the MJ11 genome. Strains were sampled during early log growth (OD_600_ ~0.25) in rich media (SWTO) prior to detectable biofilm production. Green indicated increased expression; red indicated reduced expression relative to mean expression per locus. * denoted loci for which differential expression differed at a FDR significance threshold of 0.05 (Table 4). **B)** Metabolic responses to BIOLOG compounds for-wild-type MJ11 (*binK^+^*), squid-adapted MJ11 *binK1 and* MJ11 *ΔbinK*.

**Table 5.**
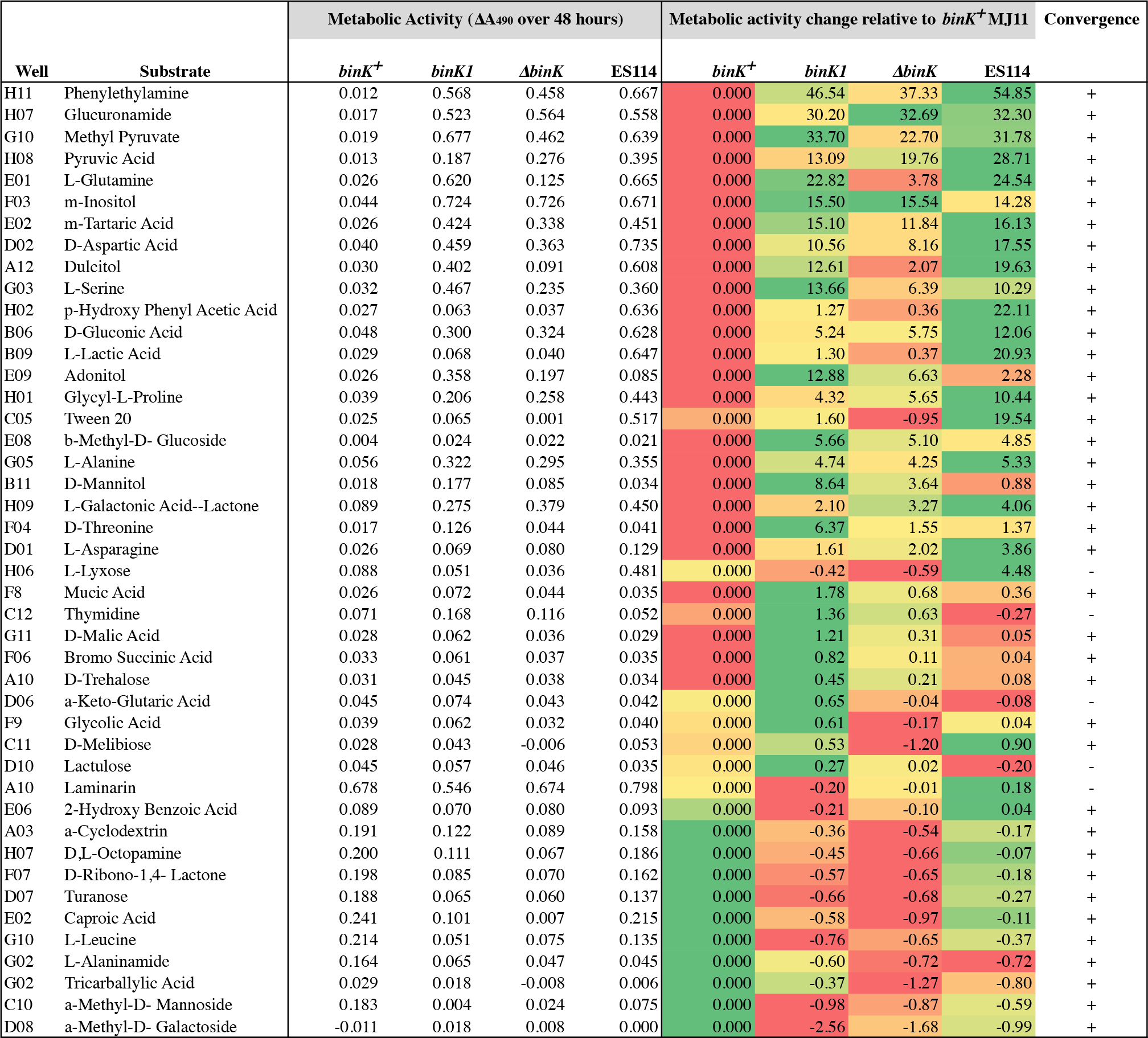
Metabolic convergence between squid native *V. fischeri* ES114 and squid-evolved *binK1*. Shown for each *V. fischeri* strain are the net changes in metabolic activity (as indicated by change in absorption of the Biolog tetrazolium redox dye) after 48 hours of exposure to each substrate. Only substrates which induced significant (FDR<0.05) differences across strains are listed. Metabolic changes in each strain relative to wild-type MJ11 *binK^+^* are colored to indicate relatively increased (green) or decreased (red) activity. Of the 190 substrates tested, 44 substrates yielded significant differences across strain, including 39 which indicate congruent metabolic responses by ES114 and *binK1* (Exact binomial test, p = 1.405e–7).

Metabolic assays corroborated that squid evolution led to convergent metabolic responses associated with light organ growth and biofilm production. *binK1* moderated activity towards compounds present either in glycans characteristic of eukaryote mucosal epithelia (Koropatkin et al. 2012) or in *Vibrio* biofilms (Table 5) (Visick 2009). Compared with MJ11, both the *binK1* and a *ΔbinK* derivative decreased metabolic activity in the presence of mannose and galactose derivatives, becoming more similar to ES114. Greater metabolism of potentially squid-provisioned chitin and amino acid derivatives by *binK* variants was also congruent with ES114 (D-glucoronic acid, L-glutamine, glucuronamide, galacturonic acid, L-glutamic acid, ß-methyl-D-glucoside)(Graf and Ruby 1998; Wier et al. 2010; Schwartzman et al. 2015). Overall, the metabolic response of *binK* variants converged significantly with ES114 (33/190 profiles; binomial test, p=0.048).

### *Squid-adapted* binK *reduces luminescence by attenuating quorum sensing*

Granting bioluminescence is the currency of this symbiosis, previous studies correlate excessive bioluminescence with poor symbiotic potential suggesting it is a likely target of host selection (Lee and Ruby 1994a; Nishiguchi et al. 1998; Visick et al. 2000). Because excessively bright luminescence by MJ11 is among the more obvious phenotypes that contrast with the relatively dim native symbiont ES114, and evidence suggests that evolved lineages converged with the native symbiont for luminescence production (Schuster et al. 2010), we evaluated the basis of reduced luminescence in *binK* variants (Figure 6). Consistent with biological assays of broth conditioned with quorum pheromone signals from in an evolved MJ11 strain harboring the *binK1* allele (Schuster et al. 2010), the evolved *binK1* allele also delayed luminescence induction (Figure 6A) suggestive of reduced priming of quorum sensing and elevated Qrr1 ncRNA accumulation (Miyashiro et al. 2010). Evaluation of *qrr1* expression using a promoter fluorescence fusion (Miyashiro et al. 2010) confirmed that *binK1* and *binK3* both increased *qrr1* expression (Figure 6), despite harboring mutations in different BinK protein domains (Figure 2C).

**Figure 6.**
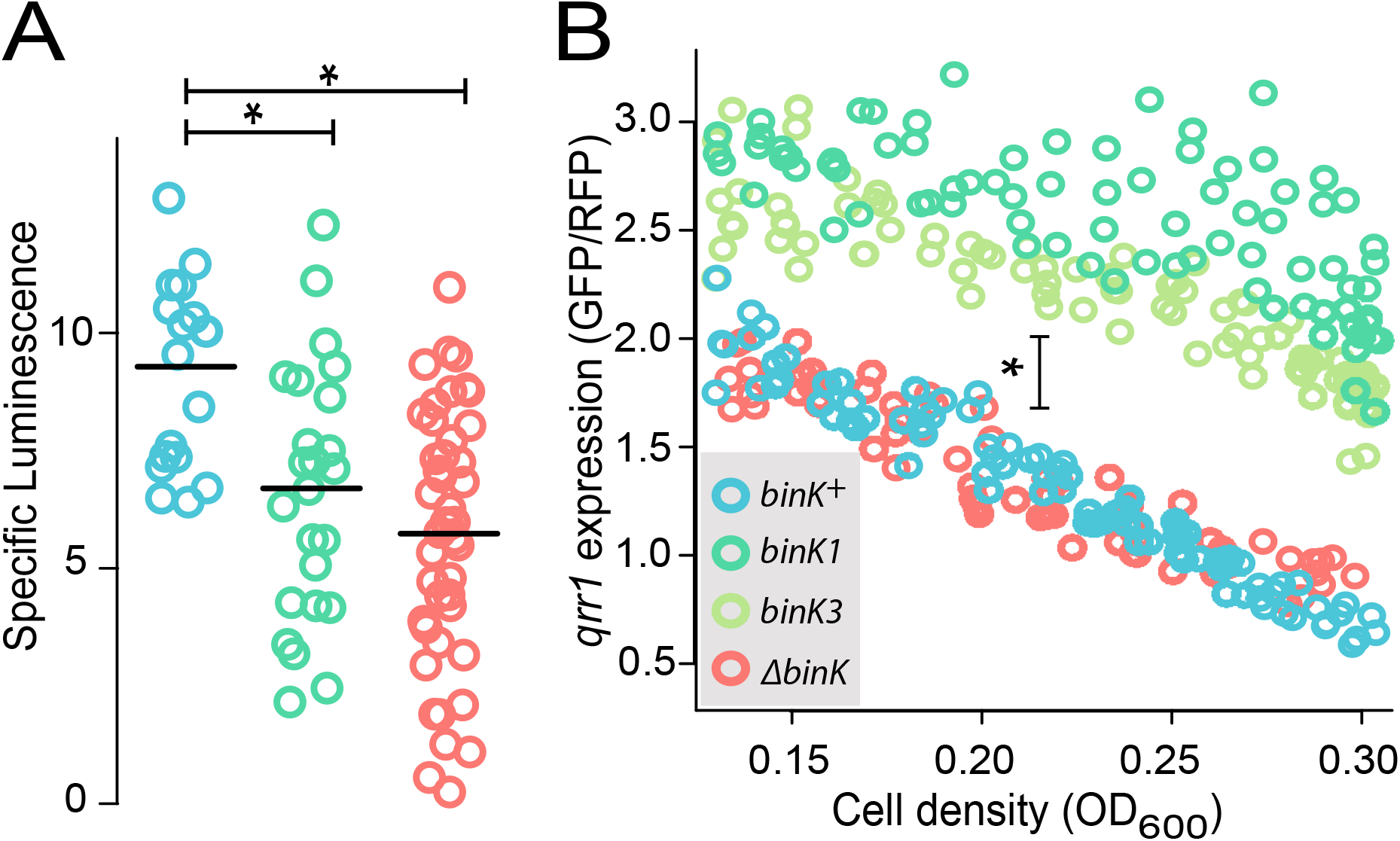
Host-adapted *binK1* attenuates luminescence. **A)** Luminescence induction at low cell density (OD_600_ < 1) as measured by relative luminescence per OD_600_. **B)** Expression of quorum-attenuator *qrr1* as measured by fluorescence of P_qrr1_-gfp relative to PtetA-mCherry (pTM268) during growth in culture. Horizontal bars show means. Error bars: std. err. *P<0.05, **P<0.01, ***P<0.001.

### *Host adaptation produced dominant, gain-of-function* binK *alleles*

Comparisons of the effects of chromosomal copies of the evolved *binK* and *ΔbinK* alleles suggested that squid selection did not favor outright loss of BinK function in MJ11, despite that the mutations mapped within and are predicted to alter the function of the protein (Figures 2, 5 & 6). The evolved *binK1* variant and the engineered *ΔbinK* mutant both improved in symbiosis traits relative to *binK*^+^ and they also did not differ significantly in their ability to produce biofilm, resist oxidative stress, evade haemocyte binding or attenuate luminesce (Figures 4 and 6A). Yet, the squid-adapted *binK* variants significantly outperformed the null mutant in colonization efficiency and in culture competition with *binK*^+^ (Figure 2D-E). Furthermore, multi-copy expression of *binK*^+^ fully restored biofilm and luminescence to wild type levels, but *binK1* only modestly rescued these traits in *ΔbinK* (Figure 4B, & 7). Together these results suggested the evolved alleles did not eliminate function. Similarly, expression of the quorum-sensing regulator *qrr1* increased significantly in *binK1* and *ΔbinK3* variants but not in the *ΔbinK* derivative (Figure 6B), suggesting a gain of function. Finally, while multi-copy expression of ancestral *binK* reduced squid colonization by all strains, multicopy expression of *binK1* enhanced colonization even in the presence of the genomic copy of the wild-type allele, demonstrating dominance of the evolved allele (Figure 7). These results indicate that evolved *binK* alleles decrease repression of biofilm traits and alter regulation of quorum sensing.

**Figure 7.**
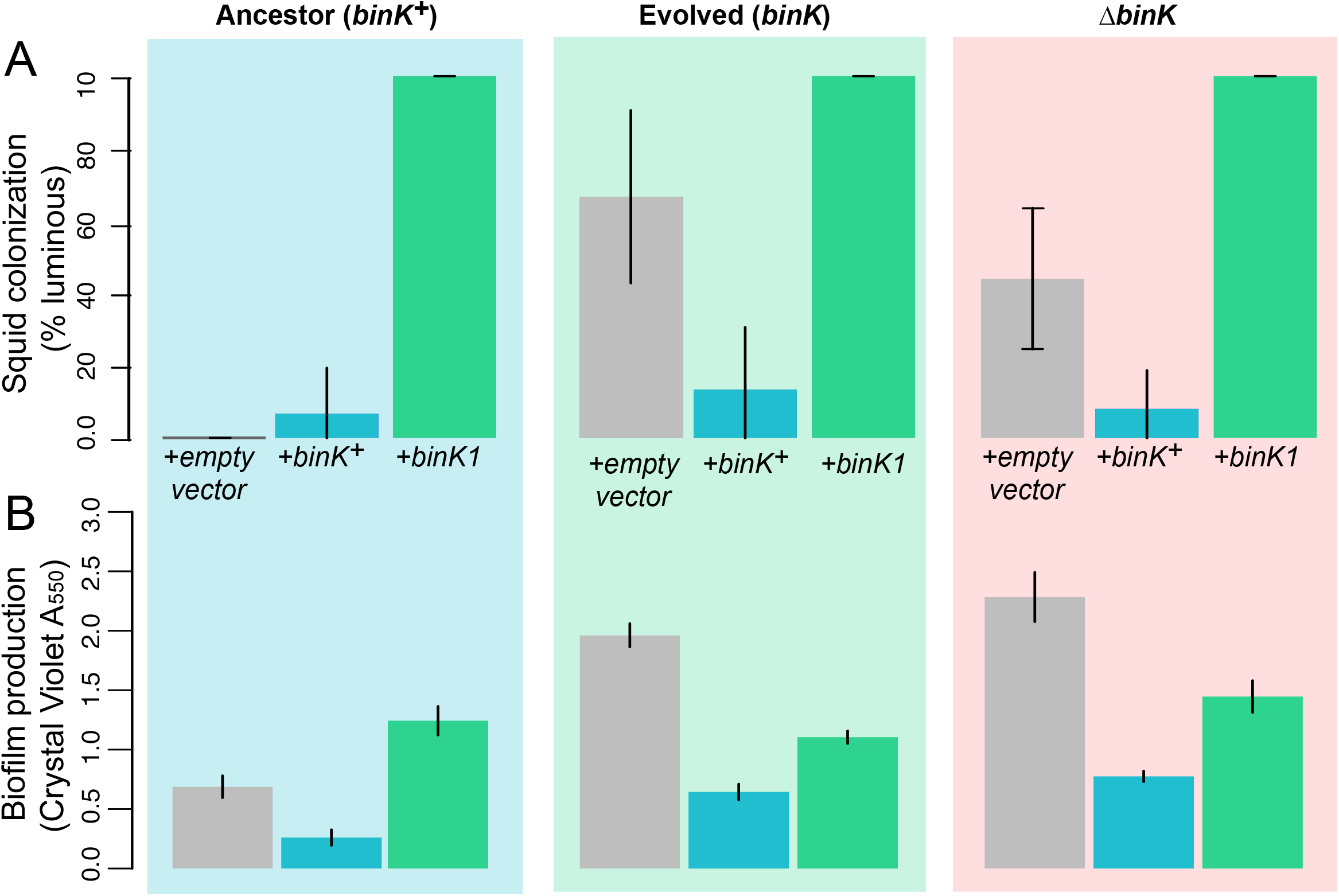
Effect of *bInK* on squid colonization, biofilm production and luminescence induction. **A)** Improvement in colonization by multicopy *in trans* expression of the evolved *binK1* allele and decreased colonization by expression of the ancestral *binK^+^* allele. Bars: 95% CI. N=15-25. **B)** Increased biofilm production by *in trans* expression of the *binK1* allele, and decreased biofilm production by expression of the ancestral *binK*^+^. Comparisons of biofilm production in control-plasmids (pVSV105) to multi-copy plasmids carrying *binK* suggest an inhibitory role of BinK in biofilm production, presumably alleviated by the dominance of the *binK1* allele. Bars: 95% CI. N=7–8. c, Luminescence induction for MJ11 strains harboring multicopy *binK*K^+^, *binK1*, or control plasmid pVSV105. Boxes designate which portion of the luminescence curves were subjected to ANOVA. A marginally significant increase in induction is seen in the mutant *ΔbinK* when complemented with *binK*^+^ (p=0.058).

## Discussion

In theory, the large population sizes and genetic diversity within bacterial species may enable symbiotic lifestyles with eukaryotic hosts to rapidly evolve. While the processes leading to pathogen emergence have been intensely studied, much less is known regarding the genetic changes that drive adaptation to novel host niches in nonpathogenic bacteria (Ochman and Moran 2001; Guan et al. 2013; Kwong and Moran 2015; Jansen et al. 2015). In pathogens, mobile elements encoded on pathogenicity islands are often cited as the cause of repeated and rapid evolution of host associations, but these elements alone rarely provide bacteria the ability to colonize hosts (Reuter et al. 2014). Further, the selective pressures exerted by new hosts may require synchronized phenotypic changes, limiting the number of adaptive ‘solutions’ available to a microbial genome constrained by regulatory structure. Here, rapid adaptation to squid symbiosis occurred in multiple parallel experimental lineages through convergent mutations in the *binK* sensor kinase gene that conferred multiple gains of function, suggesting that that the regulatory circuits of *V. fischeri* may have been pre-wired to coordinate symbiotic traits.

The convergent paths to adaptation taken by parallel evolving lineages reveals that squid hosts exert hard selection on colonizing bacteria that can produce rapid and efficacious solutions in certain unfit populations. Modelling of the evolutionary dynamics suggested that in order to survive extinction during the host-imposed bottlenecks punctuating colonization, *binK* alleles had to confer a massive fitness advantage in symbiotic association and arise early in the population expansion rather than later during symbiotic maintenance (Figures 3A & C). This prediction is consistent with the improved initiation capacity of evolved variants (Figure 4) and explains their detection in early squid passages (Table 4). Even with a theoretical accumulation of up to 185 neutral *binK* locus mutations during inoculum growth, it required an astonishing selective advantage for nine different beneficial alleles to escape extinction in half of the inoculated squid of two populations derived from different ancestors. While this could imply that an underlying mechanism promoted accumulation of mutations at this locus as a ‘hot spot’, no mutations were detected in this locus following culture evolutions of MJ11 or in ES114 (Figure 2A, Table 4)(Sung et al. 2016). Nor would these mutants be expected to rise in frequency to the point of detection considering alleles that confer enhanced fitness in squid often carried a fitness tradeoff during growth in broth culture (Figure 2E). However, the improved survival of spontaneous *qrr1*-enhancing *luxO* mutants during stationary growth suggests a plausible mechanism for ancestral population enrichment of binK alleles that enhance *qrr1* expression (Figure 2E) (Miyashiro et al. 2010; Kimbrough and Stabb 2015).

Whereas the initial culture growth of the inoculum provided genetic diversity, the key to the success of *binK* variants was only realized when under squid host selection. Estimated selective coefficients for the *binK1* allele of MJ11 ranged as high as s = 5.3 determined empirically, similar to estimates obtained by population modelling (s ~ 6) (Figure 3). Selective coefficients above 1 are rarely reported from nature; however, these are consistent with the stringent selection on pathogens as they colonize new hosts (Bedhomme et al. 2012; Morley et al. 2015; Thurman and Barrett 2016). This striking selective advantage is also consistent with the observation that ancestral populations with lower mean fitness (such as strains MJ11 and H905) are more likely than fitter populations to make a major adaptive leap (Lenski and Travisano 1994). That is, due to their distance from optimal fitness (e.g., 100% colonization), less fit ancestors are poised to benefit more from mutations of greater selective advantage (Orr 2000; 2003; Wielgoss et al. 2013). High predicted selective advantages for *binK* variants evolved from squid-maladapted strains MJ11 and H905 support the theory that adaptation from unfit ancestors may initially proceed by large shifts as opposed to incremental changes of small effect (Wiser et al. 2013).

Given that poor aggregation during initiation is a known deficiency for ancestral MJ11 (Mandel et al. 2009), it is not surprising that alteration of cell-associated matrices evidenced by enhanced biofilm production served as an early target of host selection during evolution (Figures 4 & 5). For MJ11, poor colonization capacity has been attributed to the absence of *rscS*, a horizontally-acquired activator of Syp polysaccharide that allows the native symbiont ES114 to overcome the squid initiation barrier (Yip et al. 2006; Mandel et al. 2009). However, strain H905 is closely related to ES114, resides in squid habitat and harbors *rscS* (Supplemental Figure 2), suggesting that its colonization deficiency may stem from either additional regulatory constraints on Syp production or attributes relating to a planktonic lifestyle which impair its ability to access squid light organs (Lee and Ruby 1994a). For the evolving MJ11 populations, individuals that obtained the squid-adaptive *binK1* allele dramatically improved in biofilm production and aggregation behavior at the entrance to the host light organ, thereby finding a novel regulatory solution to the initiation barrier (Figure 4). Multi-copy *sypE* repression of biofilm and impairment of initiation is consistent with Syp-mediation of improved symbiosis for the evolved variants (Figure 4), but other matrix components influenced by *binK* could also or alternatively contribute. For instance, the *binK1* allele increased expression of glycosyltransferases and sugar transporters not known to be related to Syp and wild-type *binK* repressed cellulose (Table 4 and Figure 5), suggesting that the role of *binK1* may extend beyond Syp production to alter the composition of the extracellular matrix. Even with the documented improvement in aggregation behavior (Figure 4) (Brooks and Mandel 2016), it seems unlikely that a modified extracellular matrix alone, acting as a ‘public good’ that could aid in the co-colonization of unmodified competitors, would provide sufficient fitness advantage for domination by *binK* variants.

The enhanced extracellular matrix conferred by *binK1* allele also improved survival to host defenses further promoting the success of variants in reaching the nutrient rich squid light organ crypts. Symbiotic microbes commonly secrete exopolysaccharides or glycosylated compounds to produce biofilm capsules that confer protection from macrophages, antibiotics or toxic substances, and promote adhesion to epithelial surfaces (Hsieh et al. 2003; Nizet and Esko 2009; Sengupta et al. 2013; Williams et al. 2013). Yet beyond its role in aggregate formation, it was not known whether Syp promoted squid colonization through similar mechanisms. The *binK1* allele conferred immune evasion by reducing attachment of host macrophage-like haemocytes to a level comparable with the squid-native strain ES114 and by enhancing survival to oxidative stress (Figure 4). As with biofilm production, both of these traits were suppressed by *sypE* overexpression suggesting they are Syp-mediated (Figure 4). Whereas squid immune response is mitigated by *V. fischeri* lipopolysaccharide and cell surface components (Nyholm et al. 2009) (Foster et al. 2000; Koropatnick et al. 2004; Koropatnick and Kimbell 2007), ours is the first evidence that Syp polysaccharide contributes to host immunomodulation by *V. fischeri*. Despite sharing little similarity with the capsular polysaccharide genes common to immunomodulating *Vibrio* species and other pathogens (Yildiz and Visick 2009; Shibata et al. 2012), the Syp polysaccharide gene cluster in *V. fischeri* may serve a role analogous to the polysaccharide ligands of mammalian macrophage receptors produced by gut symbionts, which also exhibit immunosuppressive activity to reduce host inflammatory response (Mazmanian et al. 2008; Chu and Mazmanian 2013; Jones et al. 2014). Recent evidence in *Vibrio parahaemolyticus* suggests that use of Syp is potentially widespread among host-associated *Vibrio*, mediating virulence and epithelial colonization (Ye et al. 2014) as well as evasion of host innate immunity (Hsieh et al. 2003; Vuong et al. 2004). The pleiotropic effects of Syp on symbiotic competence support the striking selective advantage in initiation conferred by *binK*. Further, they reveal a critical role for cell-associated polysaccharides in the squid*-Vibrio* interaction mediating not only group behaviors, but also partner selection on individual cells.

The acquisition of early light-organ colonization traits is supported by empirical estimates of enhanced fitness during the first 24 hours of colonization (Figures 3 and 4), but co-colonization studies indicated that the entire advantage associated with evolved *binK* alleles is not strictly due to improved capacity to surmount the requisite initiation bottleneck. Rather, the substantial fitness gain within hosts between 24 and 48 hours post-colonization (Fig. 3C) demonstrated that squid-adaptation through *binK* was manifested across multiple aspects of symbiosis, including traits beyond the known effects of Syp biofilm (Visick 2009; Brooks and Mandel 2016). During the maintenance stage of association (Figure 3) squid selection targets symbiont luminescence, including its intensity, and resource utilization (Graf and Ruby 1998; Visick et al. 2000; Schuster et al. 2010; Septer et al. 2013; Nishiguchi 2014; Soto et al. 2014) (Figure 1). Evolved *binK* alleles modestly attenuated and delayed the excessive luminescence by MJ11 (Figure 6), and mediated metabolic convergence with the native symbiont ES114 (Figure 5, Table 5). Concordantly, *binK* variants increased expression of the luminescence repressor and quorum-sensing regulator small RNA Qrr1 (Figure 6). These kinetic shifts support that hosts may select for dim variants from overly bright populations (Lee and Ruby 1994b), and further suggest that host selection may act not only upon luminescence but also upon the ‘private goods’ produced by key metabolic pathways that could be linked to luminescence through quorum sensing regulation (Figure 5 and Tables 4 & 5) (Dandekar et al. 2012; Davenport et al. 2015). Notwithstanding the quorum sensing regulon of metabolism by *V. fischeri* is only partially defined (Antunes et al. 2007; Qin et al. 2007; Wier et al. 2010), evolved BinK could direct metabolic restructuring through altered quorum sensing, thereby enabling nonnative colonists to quickly approximate symbiont metabolism in accordance with host mechanisms to deter cheaters (Verma and Miyashiro 2013; Davenport et al. 2015). *binK* variants also displayed patterns of increased metabolic and transcriptional activity (Figure 5, Table 2 & 5), coinciding with metabolic changes that arose in ES114 following experimental evolution in a novel host, *Euprymna tasmanica* (Soto et al. 2014). This convergence suggested responses to these metabolites could contribute to symbiont growth in juvenile squid or be co-regulated with traits directly under host selection. These metabolic shifts could promote more robust growth in light organs relative to ancestral MJ11 and account for sustained selective advantage following initial colonization (Figure 3B), but importantly these changes do not promote a general vigor phenotype (Figure 2E).

Successful symbiosis requires the coordination of numerous and diverse microbial phenotypes, and yet a broad array of traits evolved rapidly here through single adaptive mutations in an existing sensory transduction protein. These changes are remarkable when viewed from both the perspectives of genetic regulation and evolutionary biology. During bacterial evolution, sensory transduction pathways, such as two-component systems (TCS), may serve as pliable targets of selection due to the modularity of their individual components (Vogel et al. 2004; Pasek et al. 2006). The conserved phosphorelay domains (i.e. HisKA, HATPaseC, REC, HPT) and accessory domains (e.g. HAMP, CACHE) (Fig. 2C) are shared across pathways and facilitate flexible partner interaction, known as ‘cross-talk’ (Capra and Laub 2012). Signaling can evolve through limiting or expanding partnerships and output responses and is shaped by ongoing adaptation to environmental cues. Acting as environmental sensors, TCS sensor histidine kinases are effective targets of selective regimes that can act on an array of phenotypes (Bretl et al. 2011; Chambonnier et al. 2016). As such, sensor kinases can enable rapid, coordinated adaptation in part due to dual kinase and phosphatase capabilities, and ability to augment interaction with binding partners through interaction of shared modules (Capra and Laub 2012; Rowland and Deeds 2014). The array of phenotypic and physiological changes effected in MJ11 *binK* variants with reduced function in biofilm repression but altered function in quorum sensing regulation (Figure 4B, 6 & 7) implied that the evolved BinK sensor kinase may participate in more than one signal transduction pathway critical to squid symbiosis (Nyholm and McFall-Ngai 2004; Yip et al. 2006; Miyashiro and Ruby 2012). BinK could regulate Syp polysaccharide and cellulose through interactions with any of a number of identified regulators containing TCS modules (e.g. SypE, SypF, and/or SypG), partnering in their phosphorelay cascades (Brooks and Mandel 2016). Similarly, evolved BinK may attenuate quorum sensing and modify luminescence and metabolism through altered interaction with components of the quorum sensing hierarchy that includes several TCS module-containing proteins (Whistler et al. 2007; Miyashiro and Ruby 2012). The evolvable and promiscuous nature of TCS proteins may explain why genes like *rscS* and *binK* appear to play such crucial roles in the evolution of *V. fischeri* as squid symbionts. Presumably, evolved BinK enacted global effects by intersecting with pre-existing TCS circuitry, shaped by varying interactions with animal hosts during *V. fischeri* evolution (Mitrophanov and Groisman 2008; Gao and Stock 2013). Further investigations into insulation of BinK-directed pathways in different strains would give better insight into whether and how environments mold these pathways in *V. fischeri*. Regardless, the conserved but malleable components to this system make them key targets of adaptive evolution generating global effects on gene regulation.

This study demonstrates that some strains of *V. fischeri* evolve by leaps in host range by single mutations of large effect. That simple point mutations in a regulator can evoke such broad consequences reveals that disparate traits important for symbiosis initiation and maintenance are already co-regulated. Such preexisting coordination is almost certainly an evolved ability, perhaps reflective of a history of selection and ‘tinkering’ while fluctuating between free-living and host-associated environments where these bacteria naturally reside (Jacob 1977; Lee and Gelembiuk 2008). The immense populations of *Vibrio* species should in theory empower natural selection to refine even subtle traits, such as the ability to adapt to uncertain conditions and regulate various traits appropriately, with remarkable precision (Sung et al. 2016). Viewed in this light, this study suggests that the exceptional adaptability of certain bacteria like *Vibrio* to form novel intimate associations with various host organisms may be possible in part due to the structure of existing regulatory pathways formed during past transient interactions. Such parsimonious reconciliation of genomic constraints with host selection pressures is likely paramount in shaping emerging symbioses.

## Materials and Methods

### Bacterial strains, plasmids, and culture conditions

Strains and plasmids are listed in Table S1. Wild-type *Vibrio fischeri* including strain MJ11 (isolated from the fish *Monocentris japonicus* (Haygood et al. 1984)) and its derivatives, as well as squid symbiont ES114 were routinely grown at 28°C in either liquid seawater-tryptone broth (SWT) or Luria Bertani broth with added salt (LBS) with shaking at 200 rpm or on LBS medium with 1.5% agar (LBS agar) (Graf et al. 1994). *Escherichia coli* strains were routinely grown in Luria-Bertani (LB) broth (Sambrook et al. 1989) or in brain heart infusion medium (Difco) at 37°C. When required, media were supplemented with antibiotics at the following concentrations: for *V. fischeri:* chloramphenicol (Ch) at 2.5 μg/ml, kanamycin (Km) 100 μg/ml and erythromycin (Em) at 5 μg/ml; for *E. coli:* Ch at 25 μg/ml, Km at 50 μg/ml, and Em at 150 μg/ml (for HI media). For maintaining selection in seawater, these antibiotics were used at half this concentration. When applicable, agar plates were supplemented with 40 mg of 5-bromo-4-chloro-3-indolyl-β-galactopyranosidase (X-gal)/ml for visualization of β-galactosidase activity. For biofilm quantification, bacteria were grown in liquid seawater-tryptone broth with added salt (SWTO)(Bose et al. 2007). To generate transcriptomic libraries, bacteria were grown in 3mL SWTO supplemented with 0.5mM N-acetyl-D-glucosamine. Bacteria were also grown in variations of HEPES minimal medium (HMM)(Ruby and Nealson 1977), a seawater-based defined minimal medium with 1x artificial sea water (ASW: 50mM MgSO_4_, 10 mM CaCl2, 300 mM NaCl, 10 mM KCl), 0.333 mM K_2_HPO_4_, 18.5 mM NH_4_Cl, 0.0144% casamino acids buffered with 10 mM Hepes with a suitable carbon source. Other buffers were substituted and additional nutrients supplemented as follows: for *In vitro* competition, the medium was supplemented with 0.53 mM glucose; for *qrr1* expression, the medium was supplemented with 32.6 mM Glycerol, and 10 μM ferrous ammonium sulfate; for siderophore assessment in reduced iron conditions (Payne 1994) the medium was buffered with 100 mM Pipes (pH 6.8), casamino acids increased to 0.3%, and supplemented with 32.6 mM glycerol; and for qualitative detection of siderophores, this medium was additionally supplemented with 1.5% Difco bacto-agar and 10% chrome azurol S-hexadecyltrimethylammonium bromide assay solution (CAS–HDTMA) (Boettcher and Ruby 1990; Payne 1994; Lee and Ruby 1994a; Graf and Ruby 2000). Plasmids were conjugated between *E. coli* and *V. fischeri* as previously described(Stabb and Ruby 2002).

### Genome sequencing and analysis

Genomic DNA was extracted from mid-log cultures grown in LBS using Promega Wizard Genomic DNA Purification Kit (Madison, WI). Genomes of *V. fischeri* strains EM17, WH1 and H905 were sequenced *de novo* using single-molecule sequencing (Pacific Biosciences) and assembled using HGAP at the Icahn School of Medicine. Gene models for de novo genomes were predicted and annotated using Prokka with strain ES114 serving as the reference(Seemann 2014). For all strains derived from experimental evolution (both squid and culture experiments), genomic libraries were prepared following modified high-throughput Nextera library construction protocol (Baym et al. 2015) and were sequenced using the Illumina Hi-Seq 2500 platform at the University of New Hampshire or the New York Genome Center. Nextera PE adapter sequences were removed from raw reads using Trimmomatic (Bolger et al. 2014). Processed reads were aligned and analyzed against their respective strain reference (ancestral) genome to identify mutations, using default settings in breseq (Deatherage and Barrick 2014) for single isolate genomes and using the ‘–polymorphism’ setting for libraries constructed from pooled isolate gDNA. On average, 99% of the processed reads from each isolate mapped to their reference genome, resulting in an average chromosomal coverage of 95x per isolate (Table S2) for MJ11. Mutations were called only for regions covered by a minimum of 20 reads. To identify which mutation calls reflected true evolutionary change as opposed to errors in the PacBio or NCBI reference genome, we compared each putative call across all genomes derived from the same ancestor. Potential mutation calls for strain ES114 were cross-referenced with known variants (Foxall et al. 2015). Any mutation calls shared amongst at least 50% of independently evolved strain genomes were assumed to reflect ancestral genotype and thus discarded. All mutations in the *binK* locus identified by breseq were subsequently confirmed by targeted PCR amplification and Sanger sequencing (UNH and GeneWiz).

### Phylogenetic relationships among *Vibrio fischeri*

Nucleotide sequence from published Vibrionaceae genomes (*Vibrio parahaemolyticus, Aliivibrio salmonicida, A. logei; V. fischeri* strains ES114, MJ11, SR5, ZF-211; Table S3) and newly generated genomes (*V. fischeri* strains H905, EM17, SA1, CG101, VLS2, PP3, WH1 WH4) were analyzed in REALPHY and RAxML to infer whole-genome maximum likelihood phylogeny under the GTRGAMMA model of nucleotide substitution(Bertels et al. 2014). Node support was estimated by running 1000 bootstrapped analyses.

### Squid colonization and experimental evolution of *V. fischeri*

Squid colonization was conducted as previously described(Whistler and Ruby 2003). Squid were routinely held in 32 ppt Instant Ocean (IO) (Blacksburg, VA) in diH_2_O water. For determining colonization efficiency, a cohort of squid were placed in bacterial inoculum derived from mid-log (OD_6_00 0.2) SWT broth cultures diluted in filtered IO. Luminescence of squid individually housed in 4 mL IO was monitored daily, and bacterial colonization was determined by plating dilutions of homogenized squid following freezing at –80°C. For starting capacity measurements, squid were exposed to inoculum for 3 hours (ES114, EM17, and WH1) or overnight (H905 and MJ11) at increasing concentrations of bacteria (from 3000–20,000 CFU per mL), until 90% of squid became colonized as determined by luminescence detection and 24 and 48 h, and direct plating of light organ homogenates at 48hr post colonization. Colonization experiments were completed with at least 10 replicate squid, included aposymbiotic control squid, and were repeated a minimum of three times.

Strains MJ11, EM17, WH1, H905, and ES114 were evolved using squid hosts as previously described (Schuster et al. 2010). Briefly, 10 aposymbiotic hatchling squid were inoculated in an ancestral population of each strain (20,000 CFU/ml in 50 ml filtered IO for H905 and MJ11, 6000 CFU/ml for WH1, and 3000 CFU/ml EM17 and ES114). Following overnight incubation, squid were isolated and rinsed in filtered IO. Squid with detectable luminescence after 48 hours served as the founder passage for each parallel replicate population. At 96 hours following initial inoculation, squid hosts were preserved at −80°C while their seawater containing ventate was used to inoculate a new passage of aposymbiotic squid. Half of the ventate was preserved by freezing in 40% glycerol at –80°C. Serial passaging with 1ml ventate combined with 1 mL fresh IO was initiated with a hatchling squid held overnight to confirm they were uncolonized based on luminescence measurements. Passaging continued in this manner for a total of 15 host squid per experimental lineage (see Figure 1C).

Isolates from various passages of the evolutions were recovered and stored from archived ventate. Ten microliters of the ventate was plated onto SWT agar, incubated at 28°C, and representative colonies that were phenotypically similar to *V. fischeri* were quadrant streaked for isolation on LBS agar. Isolated colonies were grown in LBS liquid media and preserved by freezing in 40% glycerol at –80°C for subsequent analysis. For isolates whose identity as *V. fischeri* was suspect due to morphological differences, luminescence was measured from SWT cultures, and the strain diagnostic *gapA* gene was amplified and sequenced using primers gapA F1 and gapA R1 (Table 6) for confirmation (Nishiguchi et al. 1998).

**Table 6.**
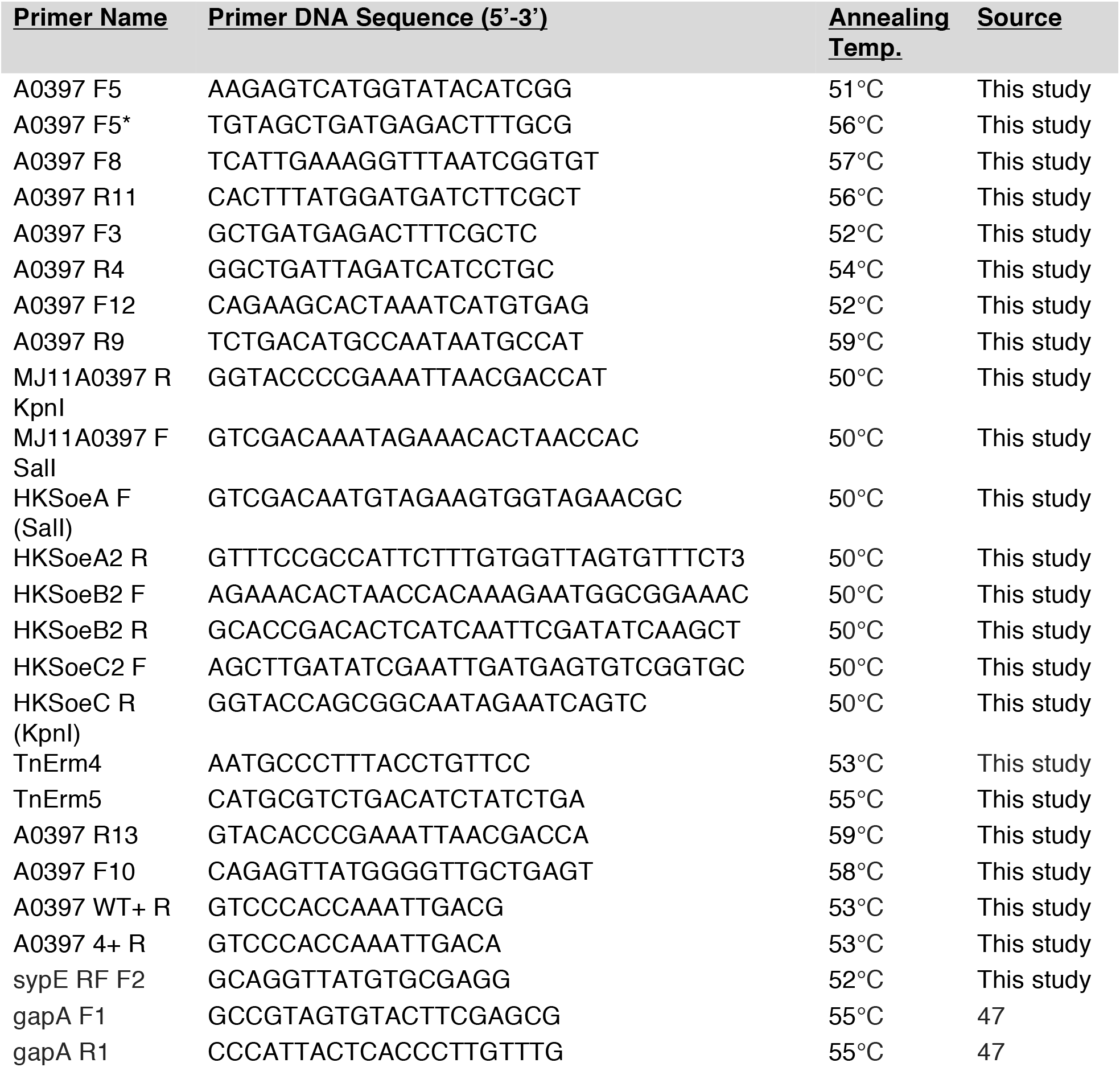
DNA Oligonucleotide primers used in this study.

### BinK orthology and hybrid histidine kinase phylogeny

To construct a gene tree for hybrid histidine kinases across *V. fischeri* strains and *Vibrio* relatives, each of the gene models from the complete genomes listed in Table S3 were queried with the PFAM Hidden Markov Models for HATPase C (PF02518), HisKA (PF00512), and REC (PF00072) domains using hmmer(HMMER n.d.). Sequences containing all of these conserved domains were then aligned in MAFFT(Katoh et al. 2002). A maximum likelihood topology was inferred using RAxML(Stamatakis 2006) under the PROTGAMMAWAG model of amino acid substitution. Gene families were annotated based on consensus among strain ES114, *Vibrio parahaemolyticus*, and *E. coli* annotations identified using the BLAST algorithm(Camacho et al. 2009).

### Recombinant DNA techniques and PCR

Integrated DNA Technologies (Coralville, IA) synthesized oligonucleotide primers listed in Table S5. Routine PCR was performed using AccuStart II PCR Supermix (Quanta, Houston, TX). Phusion High Fidelity DNA polymerase (New England Biolabs, Ipswich, MA) was used for cloning and to produce templates for sequencing reactions. PCR cycling was performed according to manufacturer’s protocol in an Eppendorf Mastercycler or Master Cycler Nexus (Eppendorf, Hamburg, Germany). Annealing temperatures used for primers were determined by subtracting 2°C from the melting temperatures (Tm) determined by Premiere Biosoft’s Netprimer. The lowest annealing temperature of the primers in the reaction was used during PCR (Table S5).

Standard molecular methods and manufacturer protocols were used for transformations, restriction enzyme digests, ligations, gel electrophoresis, and PCR. Restriction enzymes were purchased from New England Biolabs (Beverly, Massachusetts), and T4 DNA Ligase was from Invitrogen (Waltham, MA). Gel isolation and extraction of DNA from restriction digests were done using the Qiagen QIAquick Gel Extraction Kit (Qiagen, Valencia, Calif.). Plasmids for recombinant work and for sequencing were purified using Zymo Research Zyppy Plasmid Mini Prep (Irvine, CA). Genomic DNA used in PCR reactions was isolated by phenol/chloroform extraction method (Wilson 2001).

### Allele identification

Isolates from the second squid ventate from replicate MJ11 population 4 were screened for *binK* and *binK1* alleles using forward primer A0397 F5* and allele specific reverse primers A0397 WT+ R and A0397 4+ R for *binK* and *binK1* respectively (Table S5) where the presence or absence of amplicons was evaluated against controls including MJ11 (binK+), *binK1* variant MJ11EP2-4-1 and *ΔbinK* variant RF1A4. PCR amplification was conducted following denaturation at 95°C for 30 seconds followed by annealing at 53°C for 15 seconds, and elongation at 72°C for 50 seconds. To confirm identity of alleles, the *binK* region in 5 isolates was amplified by PCR using A0397 F10 and A0397 R13, and unconsumed dNTPs and primers removed using ExoSAP-IT (Affymetrix Santa Clara, CA) before Sanger-sequencing at Genewiz (Cambridge, MA) using primers A0397 F3 and A0397 R4 (Table S5). Results were aligned with reference MJ11_A0397 using Lasergene Software programs (DNASTAR, Inc. Madison, WI) and the presence of *binK1* in the evolved isolates was confirmed.

### *ΔbinK* mutant generation

The MJ11 ΔbinK::ERM (RF1A4) strain was generated by marker exchange mutagenesis using a construct produced by Splicing and Overlap Extension PCR (Horton et al. 1990). Briefly, the primer pairs HKSoeA F (SalI) and HKSoeA2 R, HKSoeB2 F and HKSoeB2 R, and HKSoeC2 F and HKSoeC R (KpnI) and the Phusion High Fidelity DNA polymerase were used to amplify the genomic region upstream and downstream of *binK* from MJ11 genomic DNA, and Em^R^ using pEVS170 plasmid DNA as the templates (Tables S1 and S5)(Lyell et al. 2008). The purified amplicons were then fused using Expand Long Template polymerase (Roche) where *binK* was replaced by an Em^R^ cassette. This purified product was cloned into pCR2.1 TOPO and transformed into TOP10 cells (Invitrogen, Waltham, MA), following the manufacturer’s protocol. Putative clones were sequenced by the Sanger method with primers M13 F, M13 R, TnErm4, and TnErm5 (Table S5) at the Hubbard Center for Genome Studies at the University of New Hampshire before the fragment was sub cloned into the suicide vector pEVS79, which was used for allelic exchange(Stabb and Ruby 2002). Whole genome re-sequencing (illumina HiSeq) confirmed the gene was replaced in MJ11 mutant RF1A4.

### Transcriptome sequencing and analysis

Single colonies of *V. fischeri* MJ11 and two of its derived strains, squid-evolved *binK1* strain (MJ11 EP2-4-1) and MJ11 mutant *ΔbinK* (RF1A4), were grown in quadruplicate until an OD_600_ of 0.25 (Biophotometer; Eppendorf AG, Hamburg, Germany) to capture populations prior to detectable biofilm activity. Cells were pelleted and flash frozen. RNA was extracted following the protocol for Quick-RNA MiniPrep kit (Zymo, Irvine, CA). Ribosomal RNA was depleted using RiboZero kit (Illumina). mRNA libraries were constructed using TruSeq Stranded mRNA library prep kit (Illumina) and sequenced using the HiSeq 2500 at New York Genome Center. Quality-trimmed reads were mapped onto the MJ11 reference genome using bowtie2(Langmead and Salzberg 2012) and quantified using RSEM(Li and Dewey 2011). Differential expression between strains was assessed using edgeR(Robinson et al. 2010) with a significance threshold of FDR < 0.05.

### Plasmid construction

*binK* and *binK1* alleles were cloned into pVSV105(Dunn et al. 2006) following amplification of MJ11 and *binK1* genomic DNA with forward primer MJ11A0397 F SalI and reverse MJ11A0397 R KpnI (Table S5). The 2.977 Kb product was cloned into pCR2.1 TOPO (Invitrogen) following the manufacturers’ instructions. The TOPO constructs were sequenced using M13F, M13R, A0397 F3, A0397 F5, A0397 F8, A0397 F12, A0397 R4, A0397 R9, and A0397 R11 (Table S5), and aligned to their respective references to ensure there were no mutations. The inserts were sub cloned from TOPO 2.1 into pVSV105 using the restriction enzymes SalI and KpnI, and ligation using Invitrogen’s T4 DNA ligase. Ligation reactions were transformed into chemically competent CC118 *λpir* cells(Herrero et al. 1990). Cell lysates of Ch^R^ colonies were directly PCR screened for insert harboring plasmids by M13F and A0397 R4. Positive clones harbored pRAD2E1(binK^+^) and pRF2A2(binK1). The *sypE* SphI and SacI fragment was sub cloned from pCLD48 into SphI and SacI digested pVSV104(Dunn et al. 2006; Hussa et al. 2008). Following transformation into chemically competent CC118 *λpir* cells, the cell lysates of Km^R^ colonies were directly screened for *sypE* insert using M13F and sypE RF F2 (Table S5). Positive clones harbored pRF2A1.

To mark bacteria for direct competition, the *lacZ* expressing plasmid pVSV103(Dunn et al. 2006) which confers a blue colony on media containing X-gal, was used along with a derivative of this plasmid (pCAW7B1) where *lacZ* was inactivated by removal of an internal 624 bp fragment by digestion with HpaI followed by self-ligation.

### Bacterial competition *in vivo*

Estimates of Malthusian growth rates and fitness for MJ11 strains were calculated by measuring relative abundances of marked strains in squid hatchings that were co-inoculated with varying ratios of each strain (Altered Starting Ratio method *sensu* (Wiser and Lenski 2015)). Strains were marked with either an intact version of the plasmid pVSV103(Dunn et al. 2006) or a version in which *lacZ* contains harboring a 200 amino acid deletion rendering LacZ unable to produce blue pigment in colonies (Table S1). Squid were inoculated overnight in 50 ml IO containing 25 μg/ml Km and stored at −80°C after 24 or 48 hours following initial inoculum exposure. To estimate CFU abundance for each strain in squid light organs, we counted blue and white colonies Kpparent after 72 hours of plating squid homogenates onto SWT plates containing 50 μg/ml Km and 1.5 mg/ml X-gal.

To calculate the selective coefficient (s) associated with the evolved variant during competition with the ancestral genotype in squid, we use the derivation in (Chevin 2011). First, Malthusian growth rates (M) were estimated by taking the natural-log of the ratio of the CFU estimate from each co-colonized light organ to the starting inoculum concentration (i.e., starting density) (Lenski et al. 1991; Lenski and Travisano 1994). Then the relative growth rate difference (*S_GR_*) was used to calculate the selection coefficient:

> Relative growth rate difference, *s_GR_* = (*M_Evo_ - M_Anc_)/M_Anc_*
>
> Selection coefficient, *s* = *s_GR_*/*In2*

Spearman rank correlation tests were then used to test for relationships between Malthusian growth rates and either starting frequency or starting density of inocula.

### Bacterial competition *in vitro*

Malthusian growth rates were estimated similarly to *in vivo* competitions where fitness for MJ11 strains was determined following co-inoculating 150 μl with a single colony from each strain marked with either pVSV103(Dunn et al. 2006) or pCAW7B1. Cultures were grown statically at 28°C and at 2 hr intervals a new culture founded by serial 1/10 dilution into fresh media. At each passage, 20 μl of each competition was diluted, and plated onto SWT plates containing 50 μg/ml Km and 1.5 mg/ml X-gal. The total number of blue and white colonies apparent after 72 hours of growth was determined and used for calculations.

### Theoretical estimation of selective advantage and mutation probability in BinK

#### Selection coefficient modelling

The analytical approximation developed in Wahl & Gerrish (2001) (Wahl and Gerrish 2001) was used to estimate the range of selection coefficients required for a novel beneficial variant to overcome the extinction risk in a population exposed to frequent bottlenecking:

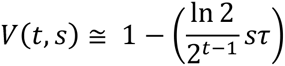

Where, *V*(*t,s*) represents the probability of extinction given selective coefficient (s) and generation (t) of growth in which the variant arises. This risk is determined by the number of generations between bottlenecks (*τ*), selective advantage (*s*), and the generation of arrival (*t*). In the context of the squid-*Vibrio* colonization dynamic, the following values were applied towards these parameters: For the initial host colonization bottleneck following inoculum growth, τ was 25 generations. For the subsequent venting bottlenecks experienced by symbiont populations, *τ* was 4 generations.

To estimate the minimum selection rate (*r*) conferred by a new rare variant capable of successfully colonizing a host (i.e, comprising one of the ~10 initiating cells (Wollenberg and Ruby 2009; Altura et al. 2013)), first the number of non-synonymous mutations to accumulate in the *binK* locus during growth of the ancestral population under neutral evolution was predicted: ~325 assuming ~25 generations of cell division to final population size of 2.4 ×10^8^. Then, using the method of Lenski & Travisano (1994)(Lenski and Travisano 1994) for estimating fitness differences in declining populations, selection rates were estimated for the rare variant using the Malthusian parameters:

*M*(rare variant) = ln(1/325)
*M*(wild-type) = ln(9/2.4 ×10^8^)
*r* = *M*(rare variant) – *M*(wild-type) = 5.6 natural logs

Using these approximations, selection coefficients for variants arising during the inoculum’s growth phase must be much larger than 1 to attain a reasonable chance of surviving the colonization bottleneck. Conversely, during the venting-regrowth periods, while the probability of a new mutation arising is low given how comparatively few generations occur during daily regrowth, beneficial alleles with coefficients as low at 0.5 may regularly survive (Figure 3C).

#### BinK mutation probability modelling

To estimate the probability of a neutral mutation occurring within the *binK* locus during either the inoculum growth phase or during growth cycles in host, the following parameters were used.

Genomic mutation rate: 2.08 ×10^−8^ bp^-1^division^-1^
Genome size of MJ11 = 4323877 bp
Available non-synonymous *binK* positions = 2595 *2/3
N_0_ Inoculum starting population = 5 cells
N_inoc_ Maximum population of inoculum prior to dilution = 2.4e8 cells
N_col_ *V. fischeri* founder population size = 12 (2-3 cells per crypt)
N_host_ Juvenile light organ *V. fischeri* population capacity = 500000 cells

Using the estimated genomic mutation rate from *V. fischeri* ES114 (Sung et al. 2016) the Poisson probability of any non-synonymous mutation in the *binK* locus approaches 1 within 19 generations of inoculum growth, while the probability of arising in the post-colonization bacterial population is below 0.001 (denoted by red line in Figure 3C).

To place the empirical observations in context of expectations using the model of Wahl and Gerrish (2001)(Wahl and Gerrish 2001), we predict that mutants carrying a selective advantage *s* ~ 2.8 would have originated within the first 10 generations of inoculum growth with the probability of any non-synonymous mutation in the locus occurring within the first 10 generations of inoculum growth to be 0.004 (under Poisson). However, the recovery of 4 distinct *binK* alleles suggests that selection could be much greater than this empirical estimation. Although quantification of the selective advantage is central to understanding the dynamics of natural selection during evolution, obtaining accurate estimates is made more difficult as fitness differentials diverge and become extreme (Wiser and Lenski 2015). We suspect that empirical estimates of *s* using competitive co-inoculations may vastly underestimate the strength of selection in this system due not only to the extreme and diverging fitness differential between ancestor and evolved strains but also to the difficulty imposed by the recovery and the challenges of accurate enumeration of rare genotypes.

### Bacterial aggregation

The capacities for MJ11 and the *binK1* variant to form cell aggregates in the squid mucus prior to entry through the ducts was conducted as previously described(Nyholm and McFall-Ngai 2003). Briefly, 1.5 hours after newly hatched squid were inoculated with ~10^5^ CFU/ml GFP-labeled strains of interest (harboring pKV111(Nyholm et al. 2000)), squid were incubated in 1 uM CellTracker Orange (Invitrogen) for 30 min, anesthetized in isotonic magnesium chloride and dissected by removing the mantel to expose the intact light organ. Dissected animals were then promptly imaged at 20X and 40X using a Zeiss laser scanning confocal microscope 510.

### Biofilm quantification

Biofilm production was quantified using a standard assay with minor modifications(OToole 2011). Briefly, a colony of bacteria from an agar plate was inoculated into either 150 μl (in a Costar 96-well plate), or 1 mL (in a 15mm glass tube) of SWTO and grown shaking at 200 rpm for 17 hours at 28°C. The biofilm that remained after expulsion of liquid, rinsing, and heat fixation at 80°C for 10 minutes was stained with 0.1% crystal violet and then decolorized in a volume of 200 μl. Biofilm production was determined by absorbance at 550 nm using a Tecan Infinite M200 plate reader.

### Host haemocyte binding of bacteria

Squid macrophage-like haemocytes were isolated from aposymbiotic hatchling squid using glass adhesion and then stained with Cell Tracker Orange (Invitrogen) suspended in Squid-Ringers prior to exposure to GFP-labeled *V. fischeri* cells following protocol previously detailed (Nyholm et al. 2009; Collins and Nyholm 2010) with modifications by Dr. Bethany Rader (personal communication). Haemocytes were exposed for one hour to *V. fischeri* strains ES114, MJ11 (*binK+*), MJ11EP2-4-1 (*binK1*) or non-symbiotic *Vibrio harveyi* B192, carrying the GFP plasmid pKV111(Nyholm et al. 2000). To test for the effect of Syp biofilm on haemocyte binding, additional assays were conducted using GFP-labeled strains carrying either control plasmid (pVSV104) or *sypE* expression plasmid (pRF2A1) in addition to GFP plasmid (pKV111)(Nyholm et al. 2000) (Table 1). Following exposure, haemocyte response to bacteria was visualized at 63x magnification by confocal microscopy and differential interference contrast using a Zeiss LSM 510. Haemocyte binding was quantified by enumeration of bound *Vibrio* relative to total *Vibrio* within a 60 μm radius surrounding each cell. Significant differences in mean proportional binding across strains were detected using a permutation-based test of independence in the R package ‘coin’.

### Hydrogen peroxide survival

Strains were grown in LBS media at 28°C with shaking at 200 rpm until cultures reach an OD600 between 1 and 1.5, the cultures were normalized to an OD_600_ of 1.0 by dilution and 5 μl was subject, in triplicate, to exposure to hydrogen peroxide at different concentrations (ranging from 0.02%-0.18%) in 200 μl of LBS media in a 96-well Costar polystyrene plate. The minimum concentrations of hydrogen peroxide that restricted all growth (MIC) of wild-type MJ11 and ES114 after over-night incubation was determined for every batch of hydrogen peroxide. Experimental concentrations ranged from 0.02%-0.18%. Differences in survival (binomial outcomes) were evaluated for significance using Fisher-Pitman permutation tests.

### Siderophore production

Siderophore was measured qualitatively as an orange halo appearing around cells cultured on CAS agar^41^ or from cell free supernatants after 17 h of growth under iron limited conditions using a chrom-azurol S liquid assay^39,74^. Colorimetric reduction in OD_630_ was measured in a Tecan Infinite M200 plate reader and % siderohpore units were calculated and normalized by cell density^39^. Siderophore units were below the detection limit for MJ11 and its *binK1* derivative but not ES114.

### Relative luminescence

At regular intervals, luminescence was quantified with a Turner 20/20 luminometer (Turner Designs, Sunnyvale, Calif.) from 100 μl of culture, which was diluted up to 1:2160 to ensure measurements were within the range of detection. The optical density of a 100 μl aliquot of *V. fischeri* MJ11 cells grown in 10 mL SWT broth culture in a 125 ml flask was determined with a Biophotometer (Eppendorf AG, Hamburg, Germany). Specific luminescence is reported as luminescence (ln) /OD_600_ of original culture. Differences in specific luminescence were tested for significance using Fisher-Pitman permutation tests on cultures sampled at low cell densities (OD_600_ < 1).

### *qrr1* expression

One colony of each MJ11 strain harboring either pTM268 (qrr1-gfp-mCherry)(Miyashiro et al. 2010) or pVSV105 (empty vector)(Dunn et al. 2006)(Table 1) grown on LBS Ch agar was suspended in 30 *μ*L minimal media. Four microliters of this suspension was inoculated into 100 *μ*L minimal medium in a flat black, clear bottom, 96 well microtiter plate (Costar). During incubation at 28°C, the OD_600_ GFP and mCherry (RFP) fluorescence was measured every hour for 45 hrs using an Infinite M200 plate reader (Tecan). GFP and RFP excitation and emission wavelengths were used as reported in (Miyashiro et al. 2010). The non-fluorescent strains harboring pVSV105 were used to set threshold fluorescence detection. Gain settings of 130 (GFP) and 160 (RFP) were determined from pilot experiments to ensure detectable fluorescence levels throughout kinetic cycles. Relative differences in GFP:RFP levels as a function of cell density (OD_600_) were evaluated between strains using Tukey’s test following an ANOVA, using a significance level of 0.05. To satisfy linearity assumptions, only data from log-phase growth were used.

### Metabolic profiling

Phenotype MicroArrays (Biolog, Hayward, CA) PM1 and PM2A were performed according to manufacturers’ protocols(Bochner et al. 2001) with few modifications for *V. fischeri* analysis, specifically including supplementation of IF-0 with 1% NaCl. Briefly, for each strain, enough inoculum for two replicate plates was prepared by recovering and mixing bacterial colonies into 16ml IF-0 to obtain a uniform suspension at OD_600_ 0.175 and mixed with dye D mixture (1:5 dilutions). PM1 and PM2A duplicate (ES114, *binK1-* and ΔbinK-variants) or triplicate (MJ11 and blank) plates were inoculated with 100 μl of suspension per well, and incubated at 28°C for 48 hours where OD_490_ was recorded by a Tecan Infinite M200 microplate reader every 4 hours to measure kinetic changes in color (redox state) of dye D. To determine which substrates elicited different kinetic responses among strains, we performed an ANOVA on OD_490_ values following normalization against the blank control values for each timed measurement. Significance of strain activity differences for any substrate was determined after correcting for multiple tests using a False Discovery Rate of 0.05. To quantify the overall significance of metabolic responses for MJ11 *binK1* and MJ11 *ΔbinK* converging with ES114 while diverging from MJ11, we used a binomial test under the null hypothesis that only 12.5% substrates should yield such a pattern across the four strains assayed (2*0.5^4^).

## Acknowledgments

We thank Richard Klobuchar, Chris Payne and the Monterey Bay Aquarium, and Deborah S. Millikan for *E. scolopes* specimens; Marcus Dillon, W. Kelley Thomas, and Robert Sebra for library preparation and genome sequencing expertise; Spencer Nyholm, Sarah McAnulty and Bethany Rader for guidance in performing haemocyte binding; Karen Visick for insightful guidance on symbiotic polysaccharide studies, strains and constructs; Matthew Neiditch, Brandon McDonald, Ashley Gagnon, Nicole Hinchey, Casey Roberts, and Sarah Martini for technical assistance; Louis Tisa, Alicia Ballock, Megan Striplin, Evan DaSilva, Feng Xu, Ashley Marcinkiewicz, Mark Mandel, Michelle Nishiguchi, William Soto, Stacia Sower, Kevin Culligan, Philip Gerrish, Caroline Turner, Todd Oakley and David Plachetzki for critical feedback and discussions. Funding was provided by the National Science Foundation (IOS-1258099) and the New Hampshire Agricultural Experiment Station through the USDA National Institute of Food and Agriculture Hatch program (Accession number 0216015). This is Scientific Contribution Number 2666.

## Contributions

MSP, RLF, IMS, VSC, and CAW designed and conceived the experiments, and performed data analysis and interpretations; LAP, BMS, and CAW conducted squid experimental evolution; MSP conducted transcriptome and bioinformatics analyses, immunological assays, empirical fitness assessment, theoretical modeling, statistical analysis, and designed graphics; RLF conducted genetic manipulations, allelic screens, sequence analysis, and mechanistic *in vitro* and *in vivo* allelic function studies; IMS performed *in vitro* quorum regulation experiments; MSP, RLF, IMS, LAP, BMS, RAD, MC, and CAW performed various experiments; MSP, RLF, IMS, VSC and CAW wrote and edited the manuscript. Genome and transcriptome short reads have been deposited in the SRA as BioProjects PRJNA316342 and PRJNA316360, respectively.

## Supplemental to Figure 2

**1.**
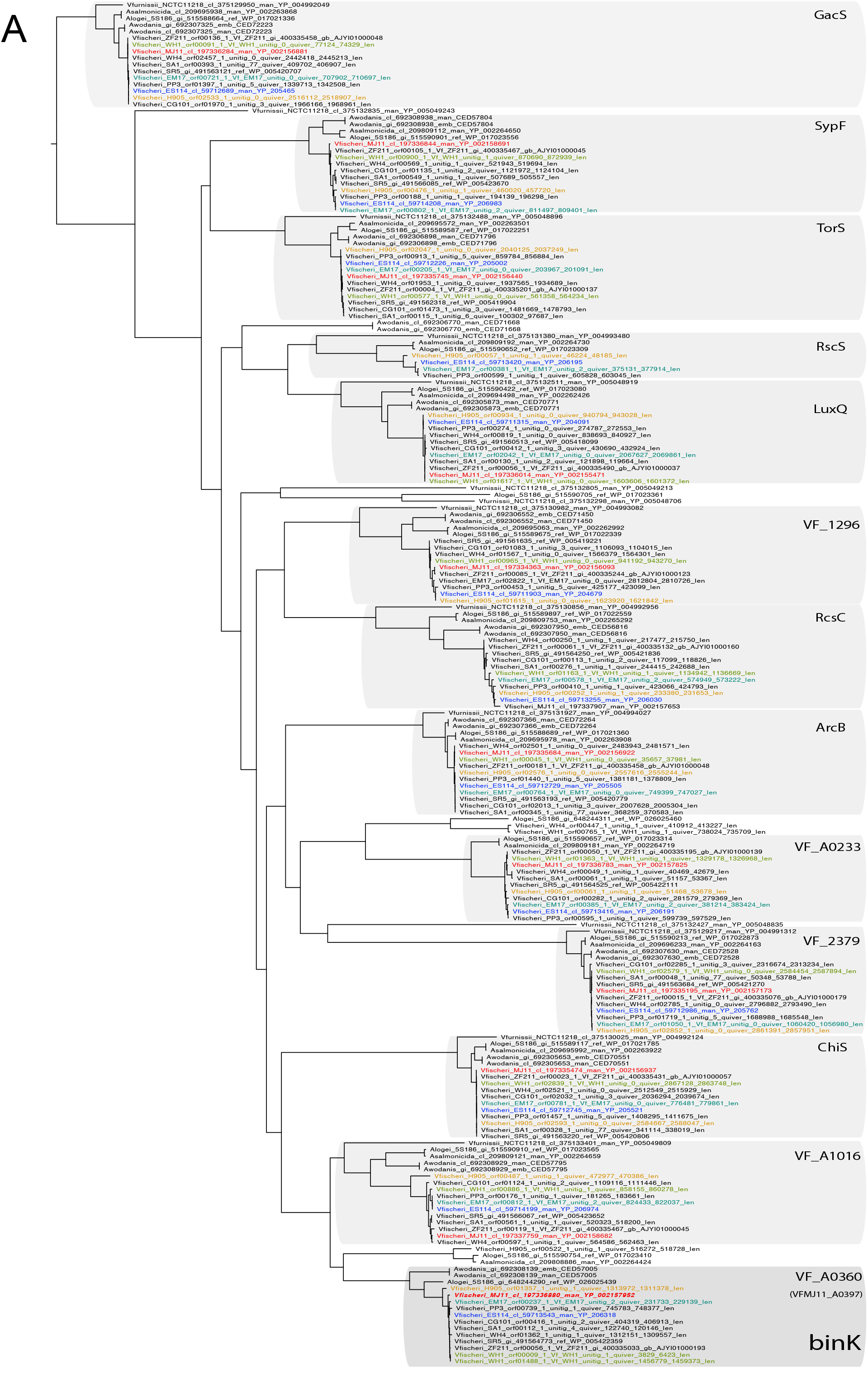
BinK orthology, conserved domains and squid-adapted *binK* alleles. Unrooted maximum-likelihood (ML) phylogeny of all hybrid histidine kinases identified in *V. fischeri* genomes. Gene families were phylogenetically annotated using *E. coli* references where possible (not shown), otherwise using the ES114 locus tag.

**2.**
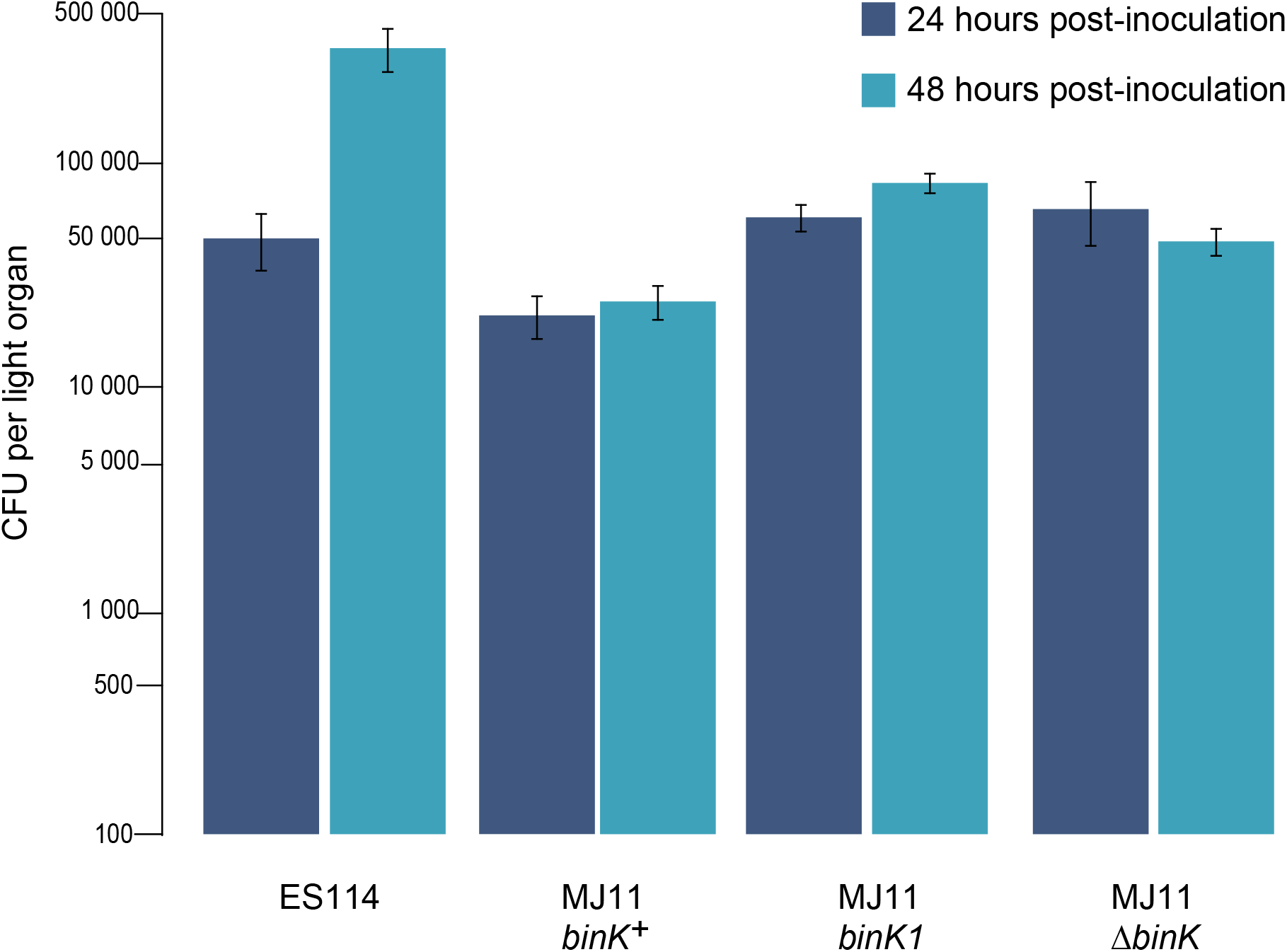
Growth of strain ES114 and MJ11 *binK* strains in squid light organs 24 or 48 hours after inoculation. Yields of symbionts determined by plating serial dilutions of squid homogenate as described previously(Whistler and Ruby 2003). Note: Y-axis is log-scaled. Bars represent standard error.

**3.**
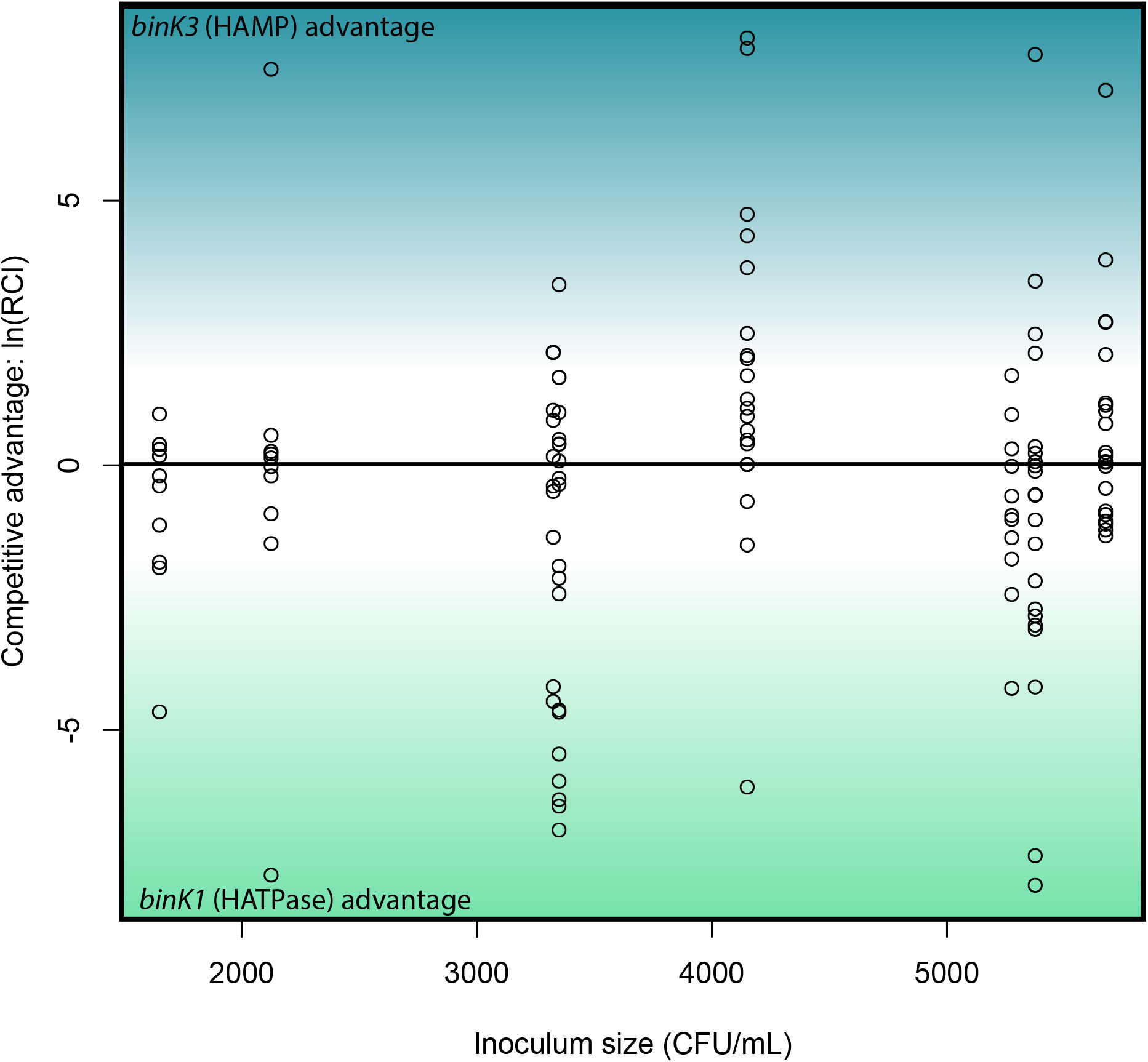
*In vivo* competitions suggest no competitive advantage to squid colonization between evolved *V. fischeri* MJ11 variants carrying either HAMP or HATPase domain mutations. Relative competitive indices for *binK1* and *binK3* MJ11 variants (carrying HATPase and HAMP domain mutations, respectively) used to co-inoculate squid across a range of inoculum densities. Points above or below zero represent squid light organs dominated by bink3 or bink1, respectively.

## Supplemental to Figure 3

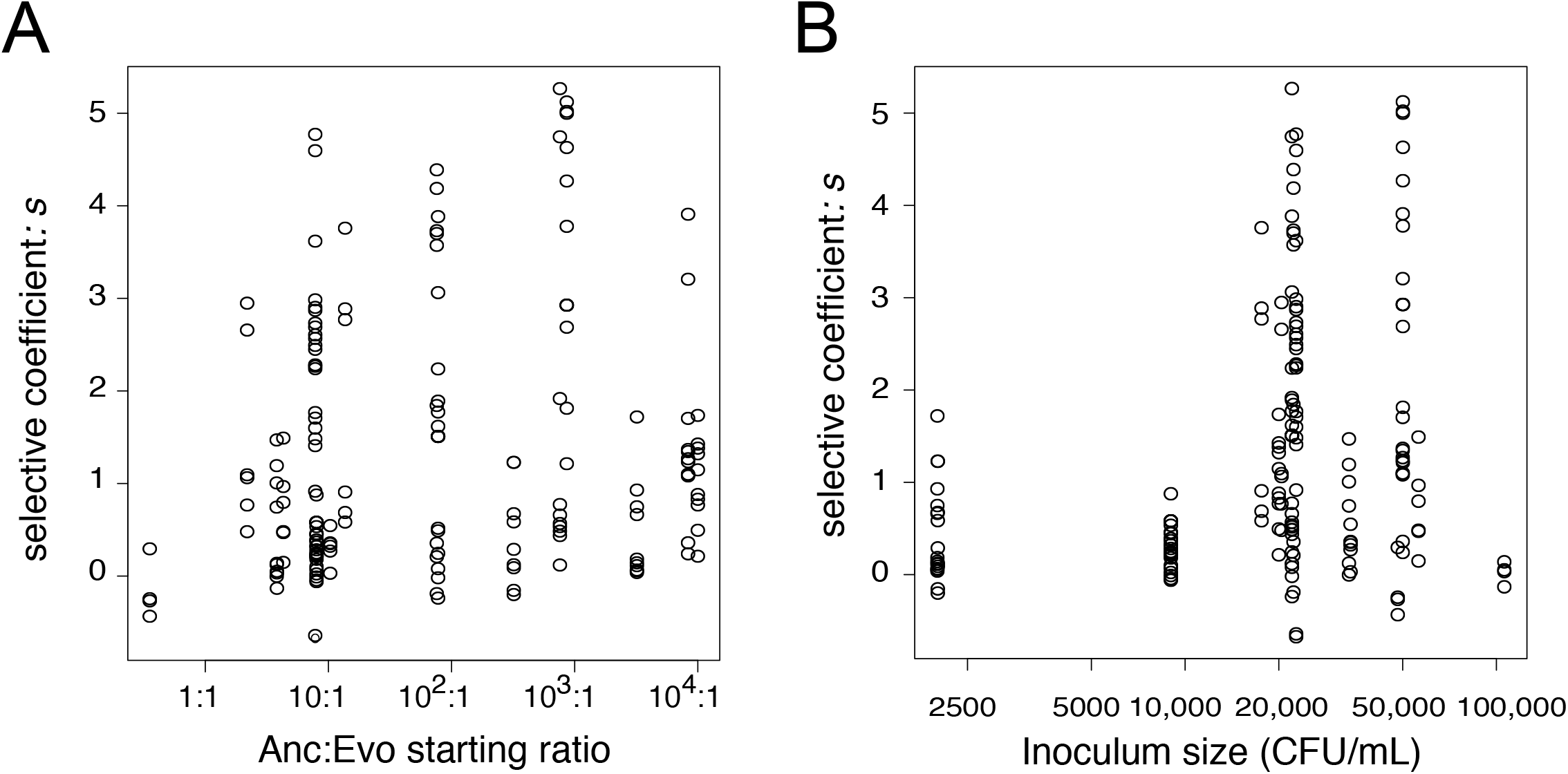
Estimated selective advantage of the evolved allele was not influenced by starting frequency (**A**) (R^2^ = 0.025, p_frequency_ = 0.62), whereas it was marginally influenced by density (**B**) (R^2^ = 0.025, p_density_ = 0.03), based on a multiple regression analysis.

## Supplemental to Figure 4

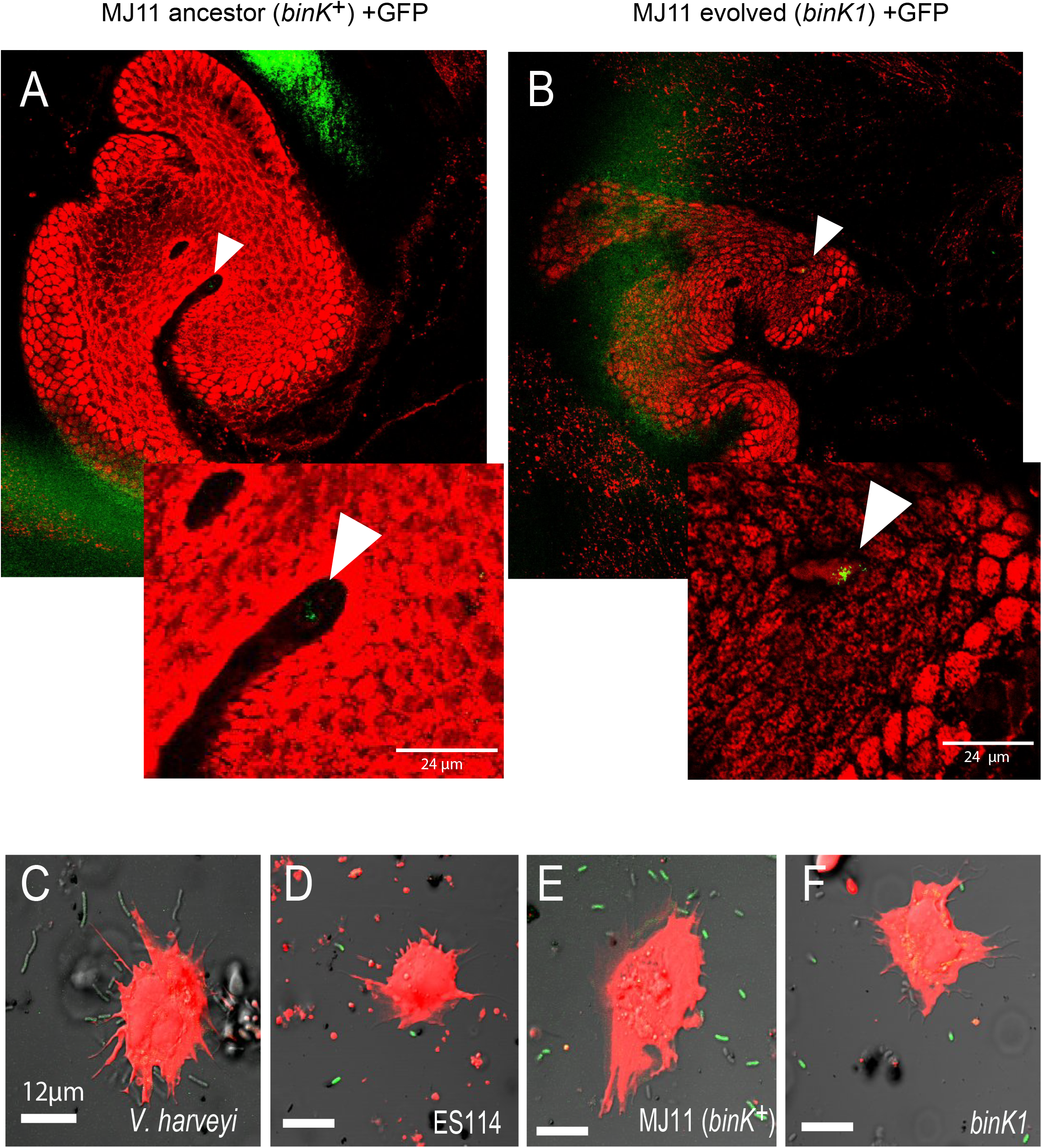
A-B) Aggregation of ancestral (A) and evolved (B) MJ11 on host mucosal epithelium prior to colonization. Host tissue stained with CellTracker Orange. Symbionts carry GFP plasmids (pKV111)(Nyholm et al. 2000). Micrographs show representative *V. fischeri* aggregates following dissection of 30 newly hatched animals incubated with each strain. Aggregates were visualized between 2 and 3 hours of inoculation using a Zeiss LSM 510 Meta laser scanning confocal microscope. **C-F) *In vitro* response of squid haemocytes to wild, squid-evolved and mutant *Vibrio* and effect of *sypE* repression of biofilm.** Micrographs show examples of haemocyte-bound non-symbiotic (*C:Vibrio harveyi*), squid-symbiotic (D: *V. fischeri* ES114), squid-naive (E: *V. fischeri* MJ11 *binK^+^*) and squid-adapted (F:MJ11 *binK1*) cells. The mean number of GFP-labelled *Vibrio* cells bound by haemocytes was quantified relative to total bacterial count in a 60 um radius using confocal microscopy at 63X magnification following one hour of bacterial exposure. Squid haemocytes in red (CellTracker Orange), *Vibrio* in green (GFP). Scale bars: 12 μm.

## Supplemental to Figure 5

**1.**
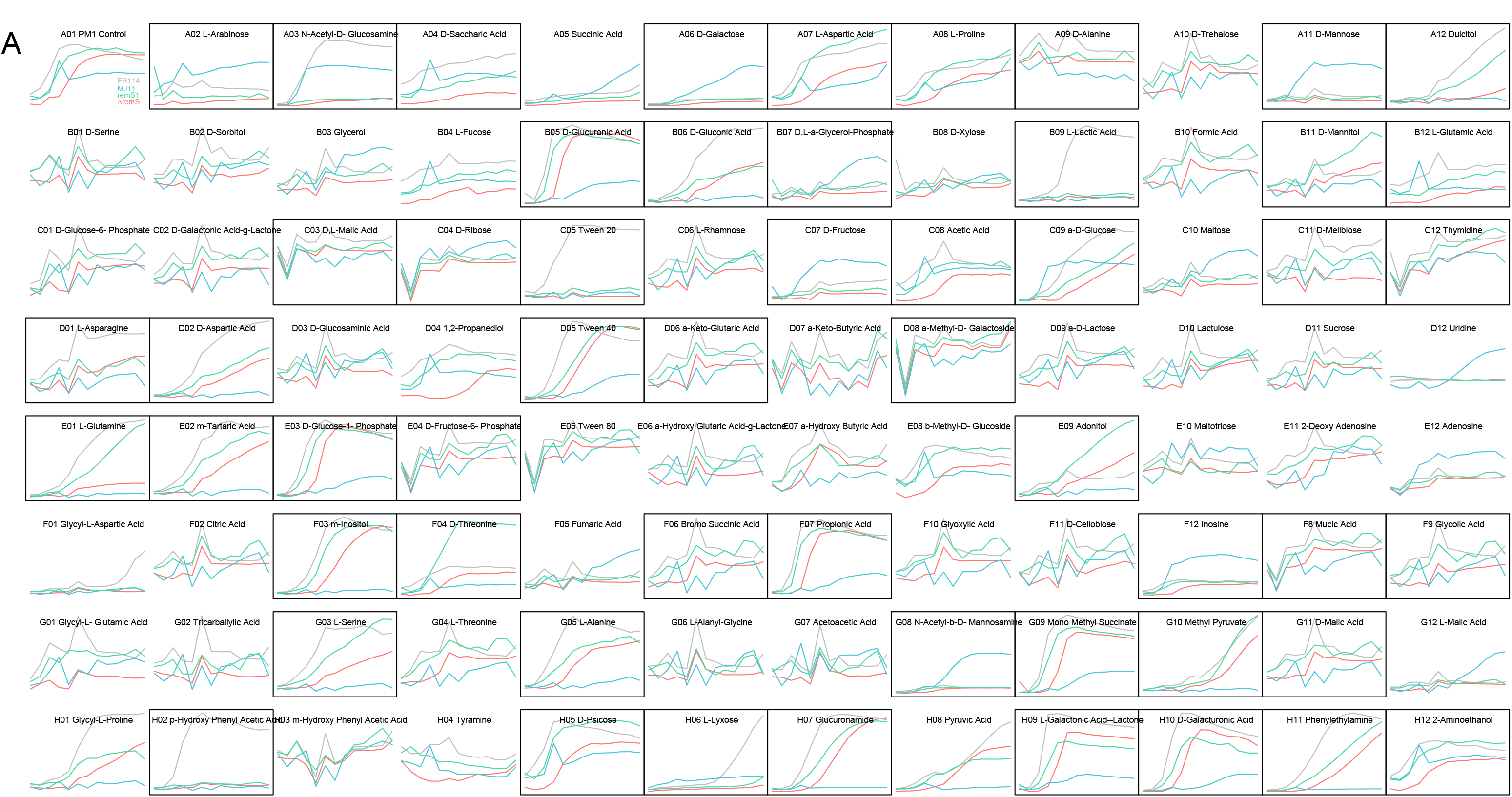
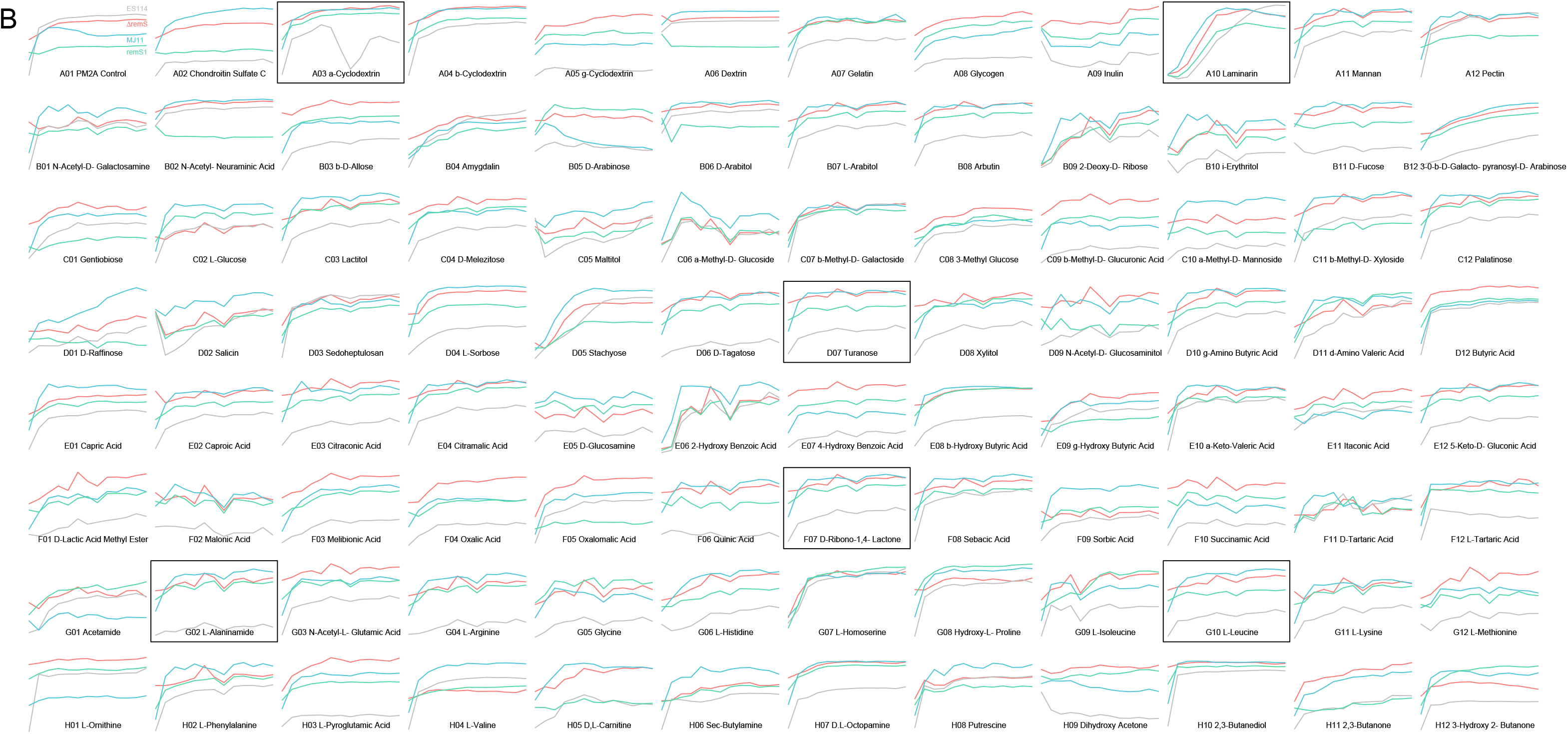
Metabolic profiles using BIOLOG phenotyping assays. Plots enclosed by boxes indicate substrates that are significantly differentially metabolized across strains (listed in Table 5). X-axis represents time (0-48 hours); Y-axis represents metabolic activity as detected by BIOLOG redox (tetrazolium) dye absorbance (OD_490_).

**2.**
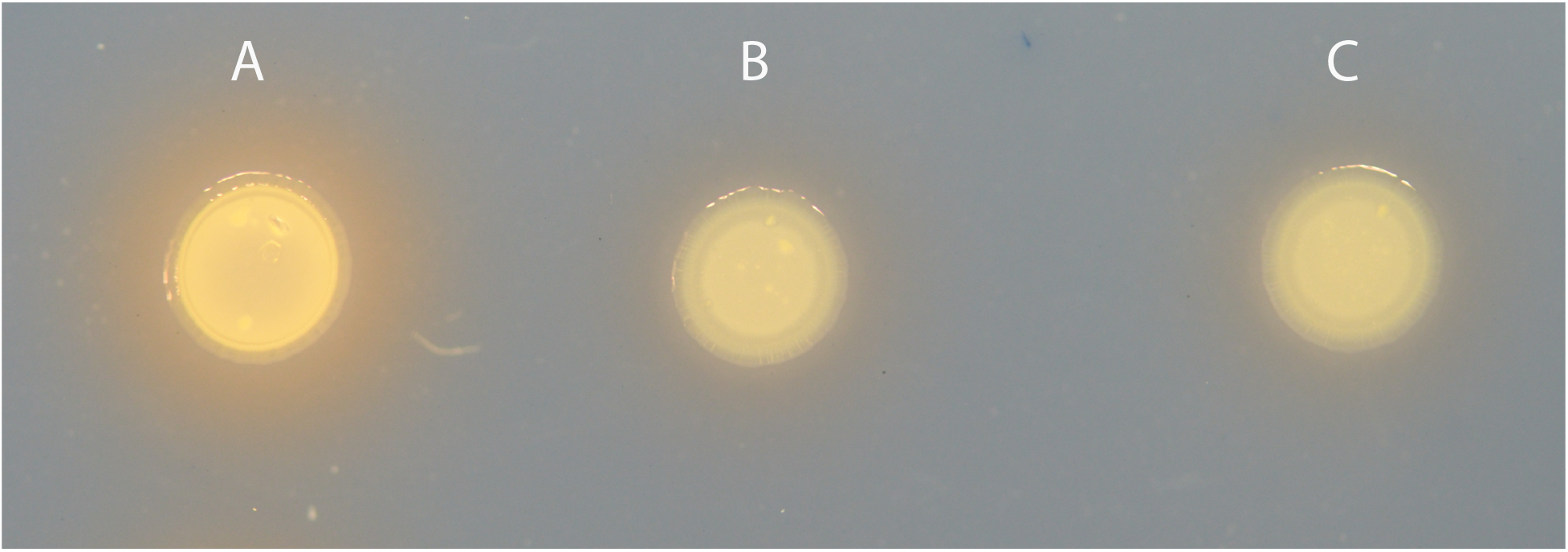
Siderophore in MJ11 and *bInK* variants.

## References

Altura, M. A., E. A. C. Heath-Heckman, A. Gillette, N. Kremer, A.-M. Krachler, C. Brennan, E. G. Ruby, K. Orth, and M. J. McFall-Ngai. 2013. The first engagement of partners in the *Euprymna scolopes-Vibrio fischeri* symbiosis is a two-step process initiated by a few environmental symbiont cells. Env Microbiol, doi: 10.1111/1462-2920.12179.

Anantharaman, V., and L. Aravind. 2000. Cache – a signaling domain common to animal Ca2+-channel subunits and a class of prokaryotic chemotaxis receptors. Trends Biochem Sci 25:535–537.

Antunes, L. C. M., A. L. Schaefer, R. B. R. Ferreira, N. Qin, A. M. Stevens, E. G. Ruby, and E. P. Greenberg. 2007. Transcriptome Analysis of the Vibrio fischeri LuxR-LuxI Regulon. J Bacteriol 189:8387–8391.

Bassis, C. M., and K. L. Visick. 2010. The Cyclic-di-GMP Phosphodiesterase BinA Negatively Regulates Cellulose-Containing Biofilms in Vibrio fischeri. J Bacteriol 192:1269–1278.

Baym, M., S. Kryazhimskiy, T. D. Lieberman, H. Chung, M. M. Desai, and R. Kishony. 2015. Inexpensive Multiplexed Library Preparation for Megabase-Sized Genomes.

Bedhomme, S., G. Lafforgue, and S. F. Elena. 2012. Multihost Experimental Evolution of a Plant RNA Virus Reveals Local Adaptation and Host-Specific Mutations. Mol Biol Evol 29:1481–1492.

Bertels, F., O. K. Silander, M. Pachkov, P. B. Rainey, and E. van Nimwegen. 2014. Automated Reconstruction of Whole-Genome Phylogenies from Short-Sequence Reads. Mol Biol Evol 31:1077–1088.

Bochner, B. R., P. Gadzinski, and E. Panomitros. 2001. Phenotype microarrays for high-throughput phenotypic testing and assay of gene function. Genome Res 11:1246–1255.

Boettcher, K. J., E. G. Ruby, and M. J. McFall-Ngai. 1996. Bioluminescence in the symbiotic squid Euprymna scolopes is controlled by a daily biological rhythm. J Comp Physiol A 179:65–73. Springer.

Boettcher, K., and E. G. Ruby. 1990. Depressed light emission by symbiotic Vibrio fischeri of the sepiolid squid Euprymna scolopes. J Bacteriol 172:3701–3706.

Bolger, A. M., M. Lohse, and B. Usadel. 2014. Trimmomatic: a flexible trimmer for Illumina sequence data. Bioinformatics 30:2114–2120.

Bose, J. L., U. Kim, W. Bartkowski, R. P. Gunsalus, A. M. Overley, N. L. Lyell, K. L. Visick, and E. V. Stabb. 2007. Bioluminescence in Vibrio fischeri is controlled by the redox-responsive regulator ArcA. Mol Microbiol 65:538–553.

Bretl, D. J., C. Demetriadou, and T. C. Zahrt. 2011. Adaptation to Environmental Stimuli within the Host: Two-Component Signal Transduction Systems of Mycobacterium tuberculosis. Microbiology and Molecular Biology Reviews 75:566–582.

Brooks, J. F., II, and M. J. Mandel. 2016. The histidine kinase BinK is a negative regulator of biofilm formation and squid colonization. J Bacteriol JB.00037–16–48.

Caley, M. J., and P. L. Munday. 2003. Growth trades off with habitat specialization. Proc Biol Sci 270:S175–S177.

Camacho, C., G. Coulouris, V. Avagyan, N. Ma, J. Papadopoulos, K. Bealer, and T. L. Madden. 2009. BLAST+: architecture and applications. BMC Bioinfo 10:421–9.

Capra, E. J., and M. T. Laub. 2012. Evolution of Two-Component Signal Transduction Systems. Annu. Rev. Microbiol. 66:325–347.

Chambonnier, G., L. Roux, D. Redelberger, F. Fadel, A. Filloux, M. Sivaneson, S. de Bentzmann, and C. Bordi. 2016. The Hybrid Histidine Kinase LadS Forms a Multicomponent Signal Transduction System with the GacS/GacA Two-Component System in Pseudomonas aeruginosa. PLoS Genet 12:e1006032–30.

Chevin, L. M. 2011. On measuring selection in experimental evolution. Biol Lett 7:210–213.

Chu, H., and S. K. Mazmanian. 2013. Innate immune recognition of the microbiota promotes host-microbial symbiosis. Nat Immunol 14:668–675.

Collins, A. J., and S. V. Nyholm. 2010. Obtaining hemocytes from the Hawaiian bobtail squid *Euprymna scolopes* and observing their adherence to symbiotic and non-symbiotic bacteria. JoVE, doi: 10.3791/1714.

Dandekar, A. A., S. Chugani, and E. P. Greenberg. 2012. Bacterial quorum sensing and metabolic incentives to cooperate. Science 338:264–266.

Darnell, C. L., E. A. Hussa, and K. L. Visick. 2008. The Putative Hybrid Sensor Kinase SypF Coordinates Biofilm Formation in *Vibrio fischeri* by Acting Upstream of Two Response Regulators, SypG and VpsR. J Bacteriol 190:4941–4950.

Davenport, P. W., J. L. Griffin, and M. Welch. 2015. Quorum Sensing Is Accompanied by Global Metabolic Changes in the Opportunistic Human Pathogen Pseudomonas aeruginosa. J Bacteriol 197:2072–2082.

Davidson, S. K., T. A. Koropatnick, R. Kossmehl, L. Sycuro, and M. J. McFall-Ngai. 2004. NO means “yes” in the squid-vibrio symbiosis: nitric oxide (NO) during the initial stages of a beneficial association. Cellular Microbiology 6:1139–1151.

Deatherage, D. E., and J. E. Barrick. 2014. Identification of Mutations in Laboratory-Evolved Microbes from Next-Generation Sequencing Data Using breseq.

Dillon, M. M., W. Sung, M. Lynch, and V. S. Cooper. 2015. The rate and molecular spectrum of spontaneous mutations in the GC-rich multichromosome genome of Burkholderia cenocepacia. Genetics, doi: 10.1534/genetics.115.176834/-/DC1.

Dunn, A. K., D. S. Millikan, D. M. Adin, J. L. Bose, and E. V. Stabb. 2006. New rfp- and pES213-Derived Tools for Analyzing Symbiotic *Vibrio fischeri* Reveal Patterns of Infection and lux Expression In Situ. Appl Env Microbiol 72:802–810.

Foster, J. S., M. A. Apicella, and M. J. McFall-Ngai. 2000. Vibrio fischeri Lipopolysaccharide Induces Developmental Apoptosis, but Not Complete Morphogenesis, of the Euprymna scolopes Symbiotic Light Organ. Developmental biology 226:242–254.

Foxall, R. L., A. E. Ballok, A. Avitabile, and C. A. Whistler. 2015. Spontaneous phenotypic suppression of GacA-defective Vibrio fischeri is achieved via mutation of csrA and ihfA. BMC Microbiol. 15:180.

Gao, R., and A. M. Stock. 2013. Evolutionary Tuning of Protein Expression Levels of a Positively Autoregulated Two-Component System. PLoS Genet 9:e1003927–10.

Graf, J., and E. G. Ruby. 1998. Host-derived amino acids support the proliferation of symbiotic bacteria. Proc Natl Acad Sci 95:1818–1822.

Graf, J., and E. G. Ruby. 2000. Novel effects of a transposon insertion in the Vibrio fischeri glnD gene: defects in iron uptake and symbiotic persistence in addition to nitrogen utilization. Mol Microbiol 37:168–179.

Graf, J., P. V. Dunlap, and E. G. Ruby. 1994. Effect of transposon-induced motility mutations on colonization of the host light organ by *Vibrio fischeri*. J Bacteriol 176:6986–6991.

Guan, S. H., C. Gris, S. E. P. Cruveiller, C. E. C. Pouzet, L. Tasse, A. E. L. Leru, A. Maillard, C. M. E. digue, J. Batut, C. Masson-Boivin, and D. Capela. 2013. Experimental evolution of nodule intracellular infection in legume symbionts. ISME J 7:1367–1377. Nature Publishing Group.

Guerrero-Ferreira, R. C., and M. K. Nishiguchi. 2007. Biodiversity among luminescent symbionts from squid of the genera Uroteuthis, Loliolus and Euprymna (Mollusca: Cephalopoda). Cladistics 23:497–506.

Haygood, M. G., B. M. Tebo, and K. H. Nealson. 1984. Luminous bacteria of a monocentrid fish *(Monocentris japonicus)* and two anomalopid fishes *(Photoblepharon palpebratus* and *Kryptophanaron alfredi):* population sizes and growth within the light organs, and rates of release into the seawater. Mar Biol 78:249–254.

Heath-Heckman, E. A. C., and M.J. McFall-Ngai. 2011. The occurrence of chitin in the hemocytes of invertebrates. Zoology (Jena) 114:191–198.

Herrero, M., V. de Lorenzo, and K. N. Timmis. 1990. Transposon vectors containing non-antibiotic resistance selection markers for cloning and stable chromosomal insertion of foreign genes in gram-negative bacteria. J Bacteriol 172:6557–6567.

Horton, R. M., Z. L. Cai, S. N. Ho, and L. R. Pease. 1990. Gene splicing by overlap extension: tailor-made genes using the polymerase chain reaction. BioTechniques 8:528–535.

Hsieh, Y. C., S. M. Liang, W. L. Tsai, Y. H. Chen, T. Y. Liu, and C. M. Liang. 2003. Study of Capsular Polysaccharide from Vibrioparahaemolyticus. Infect. Immun. 71:3329–3336.

Hussa, E. A., C. L. Darnell, and K. L. Visick. 2008. RscS Functions Upstream of SypG To Control the syp Locus and Biofilm Formation in *Vibrio fischeri*. J Bacteriol 190:4576–4583.

Jacob, F. 1977. Evolution and tinkering. Science 196:1161–1166.

Jansen, G., L. L. Crummenerl, F. Gilbert, T. Mohr, R. Pfefferkorn, R. Thänert, P. Rosenstiel, and H. Schulenburg. 2015. Evolutionary Transition from Pathogenicity to Commensalism: Global Regulator Mutations Mediate Fitness Gains through Virulence Attenuation. Mol Biol Evol 32:2883–2896.

Jones, B., and M. K. Nishiguchi. 2004. Counterillumination in the Hawaiian bobtail squid, Euprymna scolopes Berry (Mollusca: Cephalopoda). Mar Biol 144:1151–1155.

Jones, S. E., M. L. Paynich, D. B. Kearns, and K. L. Knight. 2014. Protection from Intestinal Inflammation by Bacterial Exopolysaccharides. The Journal of Immunology 192:4813–4820.

Katoh, K., K. Misawa, K.-I. Kuma, and T. Miyata. 2002. MAFFT: a novel method for rapid multiple sequence alignment based on fast Fourier transform. Nucl Acid Res 30:3059–3066.

Kawecki, T. J., R. E. Lenski, D. Ebert, B. Hollis, I. Olivieri, and M. C. Whitlock. 2012. Experimental evolution. TREE 27:547–560. Elsevier Ltd.

Kimbrough, J. H., and E. V. Stabb. 2015. Antisocial luxO Mutants Provide a Stationary-Phase Survival Advantage in *Vibrio fischeri* ES114. J Bacteriol 198:673–687.

Koropatkin, N. M., E. A. Cameron, and E. C. Martens. 2012. How glycan metabolism shapes the human gut microbiota. Nat Rev Micro 1–30.

Koropatnick, T., and J. Kimbell. 2007. Responses of host hemocytes during the initiation of the squid-Vibrio symbiosis. The Biological Bulletin.

Koropatnick, T., J. Engle, M. A. Apicella, E. V. Stabb, W. E. Goldman, and M. J. McFall-Ngai. 2004. Microbial factor-mediated development in a host-bacterial mutualism. Science 306:1186–1188.

Kwong, W. K., and N. A. Moran. 2015. Evolution of host specialization in gut microbes: the bee gut as a model. Gut Microbes 6:214–220.

Langmead, B., and S. L. Salzberg. 2012. Fast gapped-read alignment with Bowtie 2. Nat Methods 9:357–359.

Lee, C. E., and G. W. Gelembiuk. 2008. Evolutionary origins of invasive populations. Evol Appl 1:427–448.

Lee, K. H., and E. G. Ruby. 1994a. Competition between Vibrio fischeri strains during initiation and maintenance of a light organ symbiosis. J Bacteriol 176:1985–1991.

Lee, K. H., and E. G. Ruby. 1994b. Competition between Vibrio fischeri strains during initiation and maintenance of a light organ symbiosis. J Bacteriol.

Lee, K. H., and E. G. Ruby. 1994c. Effect of the squid host on the abundance and distribution of symbiotic Vibrio fischeri in nature. Appl Env Microbiol.

Lenski, R. E., and M. Travisano. 1994. Dynamics of adaptation and diversification: a 10,000-generation experiment with bacterial populations. Proc Natl Acad Sci 91:6808–6814.

Lenski, R. E., M. R. Rose, S. C. Simpson, and S. C. Tadler. 1991. Long-Term Experimental Evolution in Escherichia coli. *I. Adaptation and Divergence During 2,000 Generations*. Am Nat 138:1315–1341.

Li, B., and C. N. Dewey. 2011. RSEM: accurate transcript quantification from RNA-Seq data with or without a reference genome. BMC Bioinfo 12:323.

Lupp, C., and E. G. Ruby. 2004. Vibrio fischeri LuxS and AinS: Comparative Study of Two Signal Synthases. J Bacteriol 186:3873–3881.

Lupp, C., M. Urbanowski, E. P. Greenberg, and E. G. Ruby. 2003. The Vibrio fischeri quorum-sensing systems ain and lux sequentially induce luminescence gene expression and are important for persistence in the squid host. Mol Microbiol 50:319–331.

Lyell, N. L., A. K. Dunn, J. L. Bose, S. L. Vescovi, and E. V. Stabb. 2008. Effective Mutagenesis of *Vibrio fischeri* by Using Hyperactive Mini-Tn5 Derivatives. Appl Env Microbiol 74:7059–7063.

Mandel, M. J., M. S. Wollenberg, E. V. Stabb, K. L. Visick, and E. G. Ruby. 2009. A single regulatory gene is sufficient to alter bacterial host range. Nature 457:215–218. Nature Publishing Group.

Mazmanian, S. K., J. L. Round, and D. L. Kasper. 2008. A microbial symbiosis factor prevents intestinal inflammatory disease. Nature 453:620–625.

Mitrophanov, A. Y., and E. A. Groisman. 2008. Signal integration in bacterial two-component regulatory systems. Genes & Development 22:2601–2611.

Miyashiro, T., and E. G. Ruby. 2012. Shedding light on bioluminescence regulation in Vibrio fischeri. Mol Microbiol 84:795–806.

Miyashiro, T., M. S. Wollenberg, X. Cao, D. Oehlert, and E. G. Ruby. 2010. A single qrr gene is necessary and sufficient for LuxO-mediated regulation in Vibrio fischeri. Mol Microbiol 77:1556–1567.

Morley, V. J., S. Y. Mendiola, and P. E. Turner. 2015. Rate of novel host invasion affects adaptability of evolving RNA virus lineages. Proc Biol Sci 282:20150801–7.

Morris, A. R., and K. L. Visick. 2012. The response regulator SypE controls biofilm formation and colonization through phosphorylation of the syp-encoded regulator SypA in *Vibrio fischeri*. Mol Microbiol 87:509–525.

Nishiguchi, M. K. 2002. Host-symbiont recognition in the environmentally transmitted sepiolid squid-Vibrio mutualism. Microb Ecol 44:10–18.

Nishiguchi, M. K. 2014. Microbial experimental evolution as a novel research approach in the Vibrionaceae and squid-Vibrio symbiosis. 1–14.

Nishiguchi, M. K., E. G. Ruby, and M.J. McFall-Ngai. 1998. Competitive Dominance among Strains of Luminous Bacteria Provides an Unusual Form of Evidence for Parallel Evolution in Sepiolid Squid-Vibrio Symbioses. Appl Env Microbiol 64:3209. Am Soc Microbiol.

Nizet, V., and J. D. Esko. 2009. Bacterial and viral infections. in A. Varki, R. D. Cummings, J. D. Esko, Esko, eds. Essentials of Glycobiology. New York.

Nyholm, S. V., and M.J. McFall-Ngai. 2003. Dominance of Vibrio fischeri in Secreted Mucus outside the Light Organ of Euprymna scolopes: the First Site of Symbiont Specificity. Appl Env Microbiol 69:3932–3937.

Nyholm, S. V., and M.J. McFall-Ngai. 1998. Sampling the light-organ microenvironment of Euprymna scolopes: description of a population of host cells in association with the bacterial symbiont Vibrio fischeri. The Biological Bulletin 195:89–97.

Nyholm, S. V., and M.J. McFall-Ngai. 2004. The winnowing: establishing the squid–vibrio symbiosis. Nat Rev Micro 2:632–642.

Nyholm, S. V., E. V. Stabb, E. G. Ruby, and M.J. McFall-Ngai. 2000. Establishment of an animal-bacterial association: recruiting symbiotic vibrios from the environment. Proc Natl Acad Sci 97:10231–10235.

Nyholm, S. V., J. J. Stewart, E. G. Ruby, and M.J. McFall-Ngai. 2009. Recognition between symbiotic *Vibrio fischeri* and the haemocytes of *Euprymna scolopes*. Env Microbiol 11:483–493.

Ochman, H., and N. A. Moran. 2001. Genes lost and genes found: evolution of bacterial pathogenesis and symbiosis. Science 292:1096–1099.

Orr, H. A. 2003. The distribution of fitness effects among beneficial mutations. Genetics 163:1519–1526.

Orr, H. A. 2000. The rate of adaptation in asexuals. Genetics.

OToole, G. A. 2011. Microtiter Dish Biofilm Formation Assay. JoVE, doi: 10.3791/2437.

Pasek, S., J. L. Risler, and P. Brezellec. 2006. Gene fusion/fission is a major contributor to evolution of multi-domain bacterial proteins. Bioinformatics (Oxford, England) 22:1418–1423.

Payne, S. M. 1994. Methods in Enzymology. Pp. 329–344 in Detection, isolation, and characterization of siderophores.

Qin, N., S. M. Callahan, P. V. Dunlap, and A. M. Stevens. 2007. Analysis of LuxR Regulon Gene Expression during Quorum Sensing in Vibrio fischeri. J Bacteriol 189:4127–4134.

Reuter, S., T. R. Connor, L. Barquist, D. Walker, T. Feltwell, S. R. Harris, M. Fookes, M. E. Hall, N. K. Petty, T. M. Fuchs, J. Corander, M. Dufour, T. Ringwood, C. Savin, C. Bouchier, L. Martin, M. Miettinen, M. Shubin, J. M. Riehm, R. Laukkanen-Ninios, L. M. Sihvonen, A. Siitonen, M. Skurnik, J. P. Falcao, H. Fukushima, H. C. Scholz, M. B. Prentice, B. W. Wren, J. Parkhill, E. Carniel, M. Achtman, A. McNally, and N. R. Thomson. 2014. Parallel independent evolution of pathogenicity within the genus Yersinia. Proc Natl Acad Sci 111:6768–6773.

Robinson, M. D., D. J. McCarthy, and G. K. Smyth. 2010. edgeR: a Bioconductor package for differential expression analysis of digital gene expression data. Bioinformatics 26:139–140.

Rowland, M. A., and E. J. Deeds. 2014. Crosstalk and the evolution of specificity in two-component signaling. Proc Natl Acad Sci 111:9325–9325.

Ruby, E. G., and K. H. Nealson. 1977. Pyruvate production and excretion by the luminous marine bacteria. Appl Env Microbiol 34:164–169.

Ruby, E. G., and M.J. McFall-Ngai. 1999. Oxygen-utilizing reactions and symbiotic colonization of the squid light organ by Vibrio fischeri. Trends in Microbiology 7:414–420.

Saccheri, I., and I. Hanski. 2006. Natural selection and population dynamics. TREE 21:341–347.

Sambrook, J., E. F. Fritsch, and T. Maniatis. 1989. Molecular Cloning: a laboratory manual. 2nd ed. Cold Spring Harbor Laboratory Press, Cold Spring Harbor, NY.

Schuster, B. M., L. A. Perry, V. S. Cooper, and C. A. Whistler. 2010. Breaking the language barrier: experimental evolution of non-native *Vibrio fischeri* in squid tailors luminescence to the host. Symbiosis 51:85–96.

Schuster, M., D. Joseph Sexton, S. P. Diggle, and E. Peter Greenberg. 2013. Acyl-Homoserine Lactone Quorum Sensing: From Evolution to Application. Annu. Rev. Microbiol. 67:43–63.

Schwartzman, J. A., E. Koch, E. A. C. Heath-Heckman, L. Zhou, N. Kremer, M. J. McFall-Ngai, and E. G. Ruby. 2015. The chemistry of negotiation: Rhythmic, glycan-driven acidification in a symbiotic conversation. Proc Natl Acad Sci 112:566–571.

Seemann, T. 2014. Prokka: rapid prokaryotic genome annotation. Bioinformatics 30:2068–2069.

Sengupta, R., E. Altermann, R. C. Anderson, W. C. McNabb, P. J. Moughan, and N. C. Roy. 2013. The Role of Cell Surface Architecture of Lactobacilli in Host-Microbe Interactions in the Gastrointestinal Tract. Mediators of Inflammation 2013:1–16.

Septer, A. N., N. L. Lyell, and E. V. Stabb. 2013. The Iron-Dependent Regulator Fur Controls Pheromone Signaling Systems and Luminescence in the Squid Symbiont Vibrio fischeri ES114. Appl Env Microbiol 79:1826–1834.

Septer, A. N., Y. Wang, E. G. Ruby, E. V. Stabb, and A. K. Dunn. 2011. The haem-uptake gene cluster in Vibrio fischeri is regulated by Fur and contributes to symbiotic colonization. Env Microbiol 13:2855–2864.

Shibata, S., E. S. Yip, K. P. Quirke, J. M. Ondrey, and K. L. Visick. 2012. Roles of the Structural Symbiosis Polysaccharide (syp) Genes in Host Colonization, Biofilm Formation, and Polysaccharide Biosynthesis in Vibrio fischeri. J Bacteriol 194:6736–6747.

Small, A., and M.J. McFall-Ngai. 1999. Halide peroxidase in tissues that interact with bacteria in the host squid Euprymna scolopes. Journal of cellular biochemistry 72.

Soto, W., F. M. Rivera, and M. K. Nishiguchi. 2014. Ecological Diversification of *Vibrio fischeri* Serially Passaged for 500 Generations in Novel Squid Host *Euprymna* tasmanica. Microb Ecol 67:700–721.

Stabb, E. V., and E. G. Ruby. 2002. RP4-based plasmids for conjugation between Escherichia coli and members of the Vibrionaceae. Method Enzymol 358:413–426.

Stamatakis, A. 2006. RAxML-VI-HPC: maximum likelihood-based phylogenetic analyses with thousands of taxa and mixed models. Bioinformatics 22:2688–2690.

Stewart, V., and L.-L. Chen. 2010. The S helix mediates signal transmission as a HAMP domain coiled-coil extension in the NarX nitrate sensor from *Escherichia coli* K-12. J Bacteriol 192:734–745.

Sung, W., M. S. Ackerman, M. M. Dillon, T. G. Platt, C. Fuqua, V. S. Cooper, and M. Lynch. 2016. Genetic Drift Limits Insertion-deletion Mutation-rate Evolution. G3.

Takemura, A. F., D. M. Chien, and M. F. Polz. 2014. Associations and dynamics of Vibrionaceae in the environment, from the genus to the population level. Front Microbiol 5:38.

Thurman, T. J., and R. D. H. Barrett. 2016. The genetic consequences of selection in natural populations. Mol Ecol n/a–n/a.

Travisano, M., and R. G. Shaw. 2013. Lost in the map. Evolution 67:305–314.

Verma, S., and T. Miyashiro. 2013. Quorum Sensing in the Squid-Vibrio Symbiosis. IJMS 14:16386–16401.

Visick, K. L. 2009. An intricate network of regulators controls biofilm formation and colonization by *Vibrio fischeri*. Mol Microbiol 74:782–789.

Visick, K. L., J. Foster, J. Doino, M. McFall-Ngai, and E. G. Ruby. 2000. *Vibrio fischeri* lux genes play an important role in colonization and development of the host light organ. J Bacteriol 182:4578–4586.

Visick, K., and M.J. McFall-Ngai. 2000. An exclusive contract: specificity in the Vibrio fischeri-Euprymna scolopes partnership. J Bacteriol.

Vogel, C., M. Bashton, N. D. Kerrison, C. Chothia, and S. A. Teichmann. 2004. Structure, function and evolution of multidomain proteins. Curr. Opin. Struct. Biol. 14:208–216.

Vuong, C., J. M. Voyich, E. R. Fischer, K. R. Braughton, A. R. Whitney, F. R. DeLeo, and M. Otto. 2004. Polysaccharide intercellular adhesin (PIA) protects Staphylococcus epidermidis against major components of the human innate immune system. Cellular Microbiology 6:269–275.

Wahl, L. M., and P. J. Gerrish. 2001. The probability that beneficial mutations are lost in populations with periodic bottlenecks. Evolution 55:2606–2610.

Waters, C. M., and B. L. Bassler. 2005. Quorum sensing: cell-to-cell communication in bacteria. Annual review of cell and developmental biology 21:319–346.

Weis, V. M., A. L. Small, and M.J. McFall-Ngai. 1996. A peroxidase related to the mammalian antimicrobial protein myeloperoxidase in the Euprymna-Vibrio mutualism. Proc Natl Acad Sci 93:13683–13688.

Whistler, C. A., and E. G. Ruby. 2003. GacA Regulates Symbiotic Colonization Traits of Vibrio fischeri and Facilitates a Beneficial Association with an Animal Host. J Bacteriol 185:7202–7212.

Whistler, C. A., T. A. Koropatnick, A. Pollack, M. J. McFall-Ngai, and E. G. Ruby. 2007. The GacA global regulator of Vibrio fischeri is required for normal host tissue responses that limit subsequent bacterial colonization. Cellular Microbiology 9:766–778.

Whitehead, N. A., A. M. Barnard, H. Slater, N. J. Simpson, and G. P. Salmond. 2001. Quorum-sensing in Gram-negative bacteria. FEMS Microbiol Rev 25:365–404.

Wielgoss, S., J. E. Barrick, O. Tenaillon, M. J. Wiser, W. J. Dittmar, S. Cruveiller, B. Chane-Woon-Ming, C. Médigue, R. E. Lenski, and D. Schneider. 2013. Mutation rate dynamics in a bacterial population reflect tension between adaptation and genetic load. Proc. Natl. Acad. Sci. U.S.A. 110:222–227.

Wier, A. M., S. V. Nyholm, M. J. Mandel, R. P. Massengo-Tiasse, A. L. Schaefer, I. Koroleva, S. Splinter-BonDurant, B. Brown, L. Manzella, E. Snir, H. Almabrazi, T. E. Scheetz, M. de F. Bonaldo, T. L. Casavant, M. B. Soares, J. E. Cronan, J. L. Reed, E. G. Ruby, and M.J. McFall-Ngai. 2010. Transcriptional patterns in both host and bacterium underlie a daily rhythm of anatomical and metabolic change in a beneficial symbiosis. Proc Natl Acad Sci 107:2259–2264.

Williams, D. W., R. P. C. Jordan, X.-Q. Wei, C. T. Alves, M. P. Wise, M. J. Wilson, and M. A. O. Lewis. 2013. Interactions of Candida albicanswith host epithelial surfaces. Journal of Oral Microbiology 5:303–8.

Wilson, K. 2001. Preparation of Genomic DNA from Bacteria. John Wiley & Sons, Inc., Hoboken, NJ, USA.

Wiser, M. J., and R. E. Lenski. 2015. A Comparison of Methods to Measure Fitness in *Escherichia coli*. PLoS ONE 10:e0126210–11.

Wiser, M. J., N. Ribeck, and R. E. Lenski. 2013. Long-term dynamics of adaptation in asexual populations. Science 342:1364–1367.

Wolfe, A. J., D. S. Millikan, J. M. Campbell, and K. L. Visick. 2004. Vibrio fischeri 54 Controls Motility, Biofilm Formation, Luminescence, and Colonization. Appl Env Microbiol 70:2520–2524.

Wollenberg, M. S., and E. G. Ruby. 2009. Population structure of *Vibrio fischeri* within the light organs of *Euprymna scolopes* squid from Two Oahu (Hawaii) populations. Appl Env Microbiol 75:193–202.

Ye, L., X. Zheng, and H. Zheng. 2014. Effect of sypQ gene on poly-N-acetylglucosamine biosynthesis in Vibrio parahaemolyticus and its role in infection process. Glycobiology 24:351–358.

Yildiz, F. H., and K. L. Visick. 2009. Vibrio biofilms: so much the same yet so different. Trends in Microbiology 17:109–118.

Yip, E. S., K. Geszvain, C. R. Deloney-Marino, and K. L. Visick. 2006. The symbiosis regulator RscS controls the syp gene locus, biofilm formation and symbiotic aggregation by *Vibrio fischeri*. Mol Microbiol 62:1586–1600. n.d. HMMER. Howard Hughes Medical Institute.

